# Physiological and Evolutionary Implications of Tetrameric Photosystem I in Cyanobacteria

**DOI:** 10.1101/544353

**Authors:** Meng Li, Alexandra Calteau, Dmitry A. Semchonok, Thomas A. Witt, Jonathan T. Nguyen, Nathalie Sassoon, Egbert J. Boekema, Julian Whitelegge, Muriel Gugger, Barry D. Bruce

**Affiliations:** Biochemistry & Cellular and Molecular Biol. Dept., University of Tennessee, Knoxville, TN 37996, USA; Bredesen Center for Interdisciplinary Research and Graduate Education, University of Tennessee, Knoxville, TN 37996, USA; LABGeM, Génomique Métabolique, Genoscope, Institut François Jacob, CEA, CNRS, Univ Evry, Université Paris-Saclay, 91057 Evry, France; Electron Microscopy Dept., Groningen Biomolecular Sciences & Biotechnology Institute, University of Groningen, 9700 AE Groningen, NLD; Collection of Cyanobacteria, Institut Pasteur, 75015 Paris, France; Pasarow Mass Spectrometry Laboratory, Geffen School of Medicine, University of California, Los Angeles, CA 90095, USA; Microbiology Dept., University of Tennessee, Knoxville, TN 37996, USA

**Keywords:** Cyanobacteria, Evolution, High Light Adaptation, Photosystem I, Tetrameric PSI

## Abstract

Photosystem I (PSI) were reported as trimeric complexes in most characterized cyanobacteria, yet monomers in plants and algae PSI. Recent reports on tetrameric PSI raised questions regarding its structural basis, physiological role, phylogenetic distribution and evolutionary significance. In this study, by examining PSI in 61 cyanobacteria, we show that tetrameric PSI, correlating with a unique *psaL* gene and genomic structure, is widespread in the heterocyst-forming cyanobacteria and their close relatives. Physiological studies on these cyanobacteria revealed that tetrameric PSI is favored under high light, with an increased content of novel PSI-bound carotenoids (myxoxanthophyll, canthaxanthan and echinenone). Together this work suggests that tetrameric PSI is an adaptation to high light, along with results showing that change in PsaL leads to trimeric PSI monomerization, supporting the hypothesis of tetrameric PSI being the evolutionary intermediate in the transition from cyanobacterial trimeric PSI to monomeric PSI in plants and algae.

## Introduction

Photosystem I (PSI) is integral in the light reactions of oxygenic photosynthesis in cyanobacteria, algae and plants (Golbeck, 2007). Upon illumination, PSI accepts an electron from plastocyanin or cytochrome c and transfers this electron to its major acceptor, ferredoxin. While algal and plant PSI have been reported to exist as mostly monomers (Ben-Shem et al., 2003; Kouril et al., 2005; Gardian et al., 2007; Veith and Buchel, 2007; Watanabe et al., 2011) and recently as dimers in spinach thylakoid membrane (Wood et al., 2018), cyanobacterial PSI has been reported as trimeric in most studies. The first study that showed this trimeric PSI structure was done in the cyanobacteria *Synechococcus* (Boekema et al., 1987). Later, trimeric PSI has been observed in a diverse range of filamentous and unicellular cyanobacteria (Shubin et al., 1992; Shubin et al., 1993; Tsiotis et al., 1995; Garczarek et al., 1998; Tucker and Sherman, 2000; Boekema et al., 2001; Bibby et al., 2003; Casella et al., 2017; MacGregor-Chatwin et al., 2017), including the most primitive known cyanobacterium, *Gloeobacter violaceus* PCC 7421 (Mangels et al., 2002). Early protein crystallography structures corroborated these findings of trimeric PSI in cyanobacteria (Almog et al., 1991; Jordan et al., 2001), such as the structure of trimeric PSI from the thermophilic cyanobacteria *Thermosynechococcus elongatus* BP-1 (Jordan et al., 2001). Due to these early seminal reports, it was assumed that PSI in all cyanobacteria was in a trimeric configuration, as opposed to the monomeric form found in plants and algae. Recently, a tetrameric form of PSI observed in the cyanobacteria *Nostoc* sp. PCC 7120 (Watanabe et al., 2011; Watanabe et al., 2014) and *Chroococcidiopsis* sp. TS-821 (TS-821) (Li et al., 2014; Semchonok et al., 2016; Shelaev et al., 2018) challenged this preconceived notion of PSI trimer being the sole oligomer in cyanobacteria. Yet, the PSI tetramer has not been viewed as a major oligomeric state in cyanobacteria. Although the discovery of tetrameric PSI in two species is now accepted, the mechanism controlling the oligomeric assembly and stability is not known. Furthermore, the physiological and evolutionary significance of this structural change has yet to be elucidated. A recent cryo-EM structure of PSI tetramer from TS-821 showed that tetrameric PSI is actually a dimer of dimers with two different interaction interfaces between monomers (Semchonok et al., 2016). This structure suggests subtle changes in the placement of the central subunit PsaL, yielding changes in helical bundling that has been implicated to be critical in the formation of PSI trimers.

In light of the recent discovery of tetrameric PSI, we extended the study to encompass a much larger and more diverse set of cyanobacteria to characterize the oligomeric state of PSI structure throughout the cyanobacterial phylum. To investigate an underlying mechanism for tetrameric PSI formation, we probe the correlation between PSI oligomeric states, PsaL sequence and genomic structure using bioinformatics and biochemistry techniques. Finally, although the physiological significance of the PSI tetramer was poorly understood in cyanobacteria, we have explored several growth conditions that may alter the formation of the tetramer. Our findings indicate that exposure to high light induces a significant increase in PSI tetramer formation, while also increasing carotenoid content. These results shed light on the role and function of higher order oligomers of PSI and provide insight into the possible cyanobacterial lineage associated with the origin of the plastid.

## Results

### PSI Oligomeric States in Cyanobacteria

To understand how PSI evolved from its cyanobacterial multimeric forms to the monomeric form in algae and plants, a comprehensive and unbiased understanding of the cyanobacterial PSI oligomeric forms is needed. This needs the investigation of the PSI structures through the diversity of cyanobacteria. Here, we have investigated the oligomeric states of PSI in 61 cyanobacteria including 34 heterocyst-forming cyanobacteria and five unicellular close relatives, as well as a set of 22 widely divergent unicellular and filamentous cyanobacteria for comparison (Dataset S1). For most heterocyst-forming cyanobacteria and their close unicellular relatives (HCR), tetrameric and/or dimeric PSI, besides the abundant monomeric PSI, was observed as the major PSI oligomers determined by BN-PAGE analysis (Figs 1, S2). Similarly, analysis of *T. elongatus* BP-1 and *Synechocystis* PCC 6803 consistently observed trimeric PSI by BN-PAGE (Figs 1a, S2). Specifically, 30 out of 34 heterocyst-forming cyanobacteria including poorly studied species such as *Spirirestis rafaelensis* UTEX B 2660 (Figs 1a, S2n), appeared to have tetrameric or dimeric PSI (Dataset S1). To verify that the altered electrophoretic mobility was truly indicative of the PSI tetramer, PSI oligomers for several species were examined using TEM, which confirmed their tetrameric structure (Figs 1b, S2). Surprisingly, a few heterocyst-forming cyanobacteria, *Anabaena inaequalis* UTEX B381, *Calothrix membranacea* UTEX B379, and *Cylindrospermum licheniforme* UTEX B2014 possessed only monomeric PSI (Fig. S2u). However, *Nostoc* sp. PCC 9335 represents the only species of cyanobacteria observed that had significant amounts of both trimeric and dimeric forms of PSI (Fig. S3a). Analysis of the unicellular cyanobacteria that are closely related to the heterocyst-forming cyanobacteria (Shih et al., 2013) including TS-821, *Synechocystis* sp. PCC 7509 and *Gloeocapsa* sp. PCC 7428 (Fig. S2a, b), exhibit more tetrameric, dimeric, and monomeric PSI than trimeric PSI. The two other species of *Chroococcidiopsis* that we tested, *Chroococcidiopsis thermalis* PCC 7203 (Fig. 1a, c) and *Chroococcidiopsis* sp. PCC 7434 (Fig. S3a), contained primarily monomeric PSI with some dimeric PSI. Collectively, these results revealed that besides monomeric PSI, tetrameric and dimeric PSI, instead of trimeric PSI, comprise the majority of PSI oligomers in the HCR. Trimeric PSI was either not detectable or only present as a minor species in this clade (Dataset S1), suggesting that there is a structural and potentially physiological basis for the propensity of dimeric/tetrameric PSI for these related cyanobacteria.

**Fig. 1.**
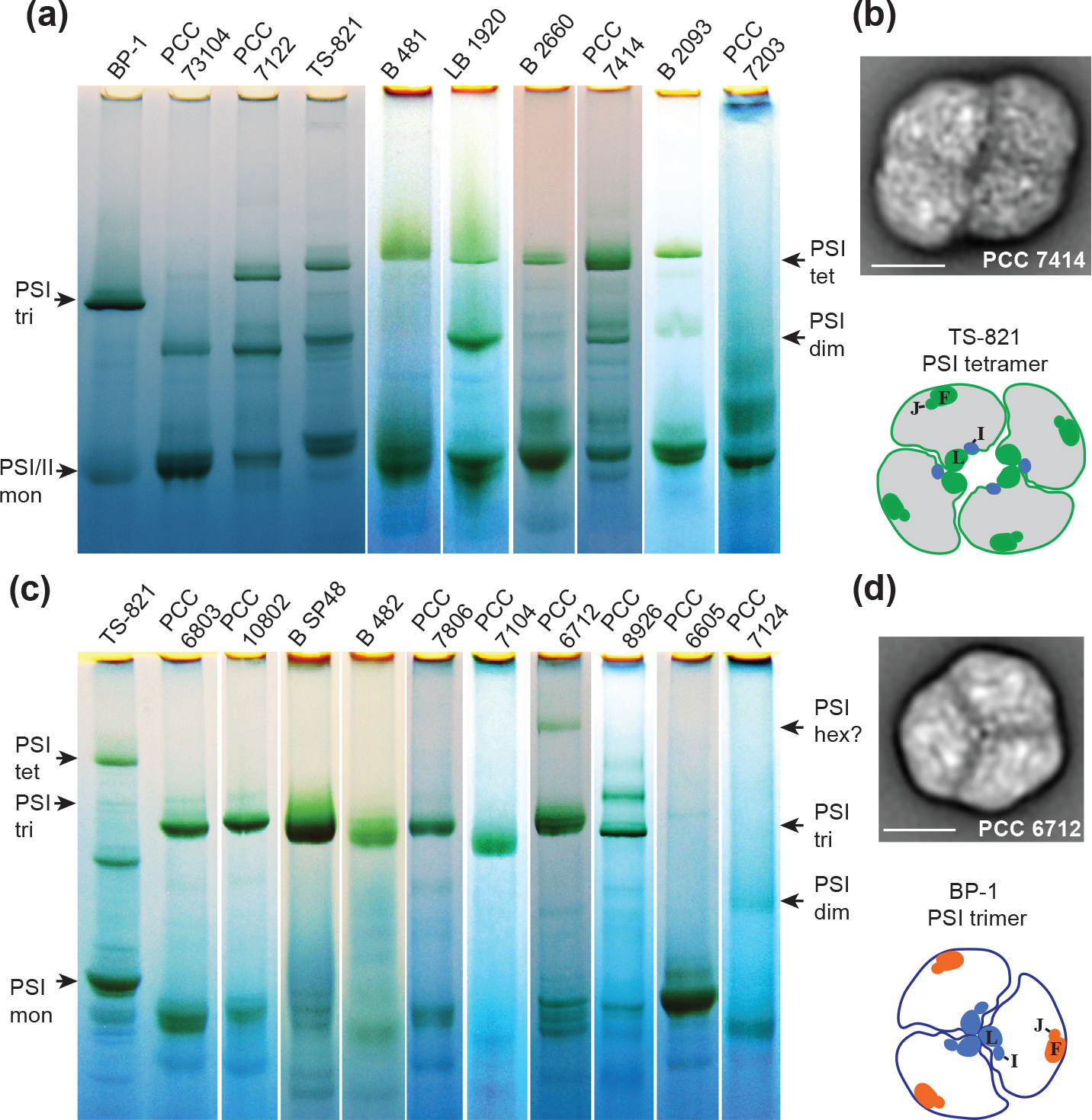
Examples of BN-PAGE and TEM analyses of PSI oligomeric states in different cyanobacteria. **(a)** BN-PAGE analyses of the PSI oligomeric states of various heterocyst-forming cyanobacteria and their close relatives. **(b)** TEM structure of PSI tetramer from PCC 7414 compared with TS-821 PSI tetramer model. **(c)** BN-PAGE analyses of the PSI oligomeric states of cyanobacteria that are distantly related to heterocyst-forming cyanobacteria. **(d)**. TEM structure of PSI trimer from PCC 6712 compared with BP-1 PSI trimer model. In **a** and **c**, for each strain, the solubilization result of 0.4 mg/mL Chl in 1% n-dodecyl-β-maltoside (DDM) is shown. TS-821, *Synechocystis* sp. PCC 6803 (PCC 6803), and *T. elongatus* (BP-1) are used as controls. The approximate migration distances for photosystem monomer (mon), dimer (dim), trimer (tri), tetramer (tet), and hexamer (hex) are indicated by arrows. The uncertainty was marked by ‘?’ symbol. In **b** and **d**, the locations of PsaL (L), PsaI (I), PsaF (F), and PsaJ (J) subunits are colored and labeled. Scale bars: 10 nm.

In contrast to HCR, most other cyanobacteria that we studied contained predominantly trimeric PSI (Figs. 1c, S2, S3; Dataset S1). When we investigated this PSI oligomer by TEM, we observed that this form was a clear trimer as shown for PCC 6712 (Fig. 1d). Amongst this group of PSI trimer-containing cyanobacteria, *T. elongatus* and *Synechocystis* PCC 6803 were included, within which the existence of trimeric PSI has been confirmed via crystallography (Jordan et al., 2001; Malavath et al., 2018). Many of the other cyanobacteria studied were also observed to contain trimeric PSI (Figs 1c, S2, S3; Dataset S1).

Occasionally, irregular PSI oligomers were observed in these diverse cyanobacteria based on results obtained via BN-PAGE analysis. For example, a potential PSI hexamer was observed in *Chroococcidiopsis* sp. PCC 6712 (Figs 1c, S2c, d) in equilibrium with the primary PSI trimer *in vitro* (Figs 1d, S2c, d). Although the BN-PAGE reveals some other minor PSI oligomeric forms in these strains, such as the PSI dimer in *Microcysti*s sp. PCC 7806 (Figs 1c, S3a, b; Dataset S1), and a potential PSI tetramer in *Planktothrix paucivesiculata* PCC 8926 (Fig. S3c), the predominant PSI oligomer in these diverse cyanobacteria is trimeric. In contrast to other PSI trimer-containing strains, the PSI monomer was clearly dominant in *Chamaesiphon minutus* PCC 6605, along with *Leptolyngbya* sp. PCC 7124, showing significant amount of PSI dimers and monomers (Figs 1c, S3a).

### Correlation of PsaL and PSI Oligomers

Previous structural studies on cyanobacterial PSI identified different PsaL-PsaL interactions within the trimeric and tetrameric PSI oligomers, shown schematically in Fig. 1b, d (Jordan et al., 2001; Semchonok et al., 2016). Previous work has identified not only subtle changes in the protein sequence of the PsaL subunits, but also a change in the genomic organization pertaining to where *psaL* is found in the cyanobacterial genome for TS-821 (Li et al., 2014). Extending this observation to the larger group of cyanobacteria presented here, we are now able to see if this correlation holds true across the different PSI oligomeric forms identified. To allow visualization of the distribution of PSI oligomers within cyanobacteria, we localized the distribution of *psaL* in relation to *psaI, psaF*, and *psaJ* genes on a species tree (Figs 2, S4a). We analyzed the *psaL* distribution of cyanobacteria with a known genomic dataset and 14 additional heterocyst-forming strains. In all HCR, we observed the *psaL* gene located downstream of *psaF* and *psaJ* (*psaF/J/L*) (Figs 2, S4b), in agreement with an earlier report (Li et al., 2014). This PsaL encoded by *psaF/J/L*, along with prevailing tetrameric/dimeric PSI, clearly delineates the clade of cyanobacteria, namely HCR, represented by green lines and the inset of Fig. 2.

**Fig. 2.**
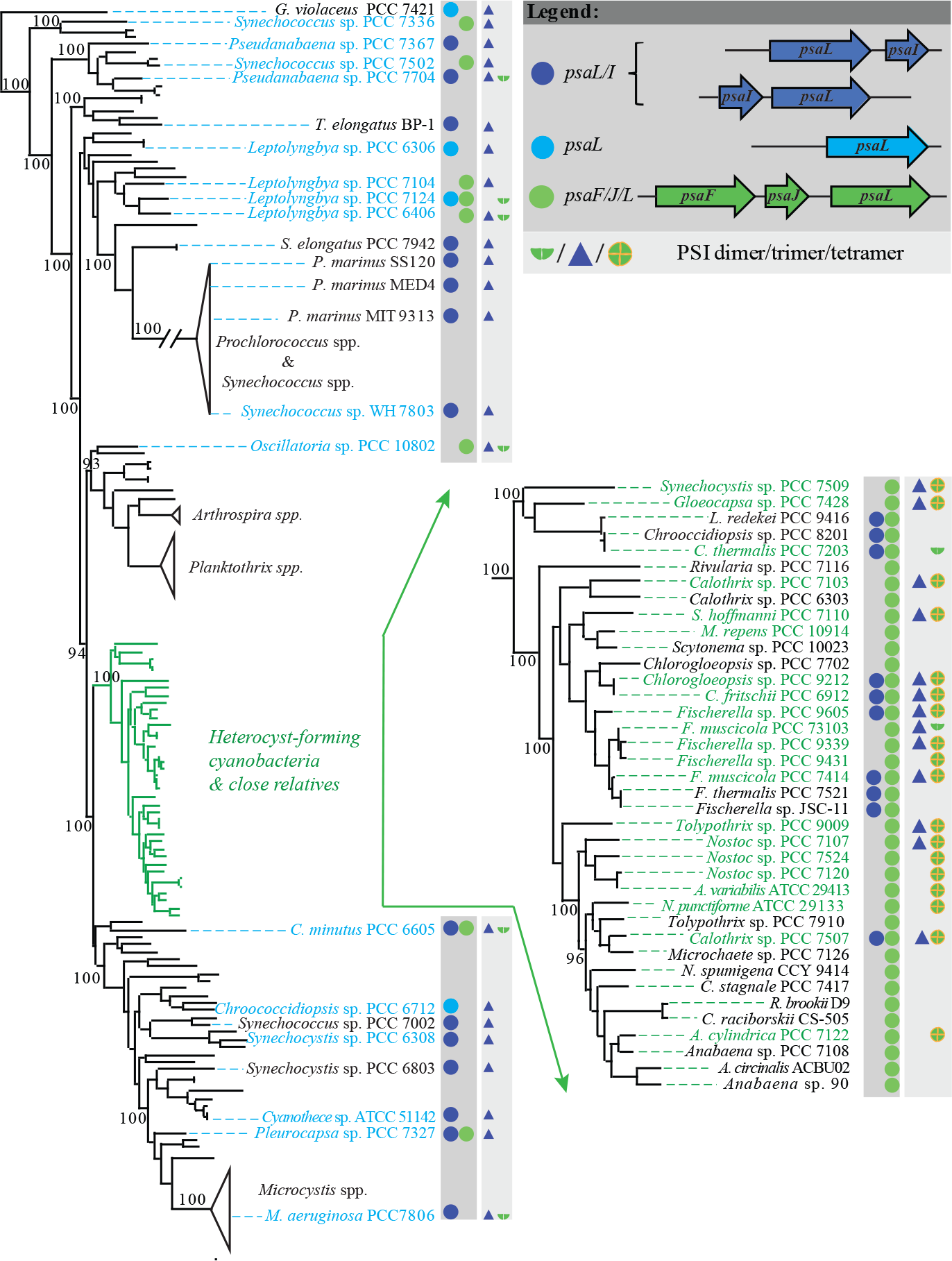
Distribution of PSI oligomers and the location of *psaL* genes among cyanobacteria with sequenced genomes. The relative location of *psaL to psaI, psaF and psaJ* genes are color coded after each strain number, followed by the observed or reported oligomeric states of PSI in that cyanobacterium. The clade including heterocyst-forming cyanobacteria and their close relatives is expanded for clarity. Bootstrap values greater than or equal to 70% are indicated by black dots. Strains whose PSI oligomeric states investigated in this study are colored in cyan or green. A fully expanded species tree is shown in Fig. S4.

Cyanobacteria that fall outside of the HCR clade often have the *psaL* flanked by *psaI* (*psaL/I*) or in a few cases found in *psaF/J/L* or isolated from other *psa* genes all together (Figs 2, S4a). Besides the 9 strains with reported PSI trimers, our analyses of the PSI oligomers identified an additional 16 strains with trimeric PSI using BN-PAGE (shown in blue text, Fig. 2). These PSI trimer-bearing cyanobacteria, spreading out in the cyanobacteria phylum outside HCR, cover each major branching point and clade. This group also includes the well characterized reference strains of *T. elongatus* and *Synechocystis* PCC 6803. When the absence of PsaL, coded by *psaF/J/L*, is clearly correlated to the PSI trimer in cyanobacteria (Fig. 2), the presence of such PsaL does not always point to a tetrameric/dimeric PSI. The phylogenetic analysis of PsaL revealed that PsaL coded by *psaF/J/L* fall in to different lineages, with the ones in HCR being a unique clade (dark green in Fig. S4b).

We did observe that some cyanobacteria with tetrameric PSI, such as *Fischerella muscicola* PCC 7414, also have two copies of the *psaL* gene, one copy organized as a *psaF/J/I* and the second as a *psaL/I* (Figs 2, S4a). The first copy (as in *psaF/J/I*) points to HCR and having tetrameric/dimeric PSI (Fig. S4b, colored in dark green) while the second form of PsaL (in the *psaL/I* locus) is closely related to the recently identified far-red light responsive PsaL, found in monomeric PSI, in *Leptolyngbya* sp. strain JSC-1 (Fig. S4b, colored in red) (Gan et al., 2014). This indicates that some of the HCR may have the second copy of the *psaL* gene (in *psaL/I*) being expressed during far-red light acclimation.

To verify a direct correlation between a specific PsaL protein and the formation of the PSI tetramer, subunit analyses for the isolated PSI trimer and tetramer from three different cyanobacteria, TS-821, PCC 7428, and PCC 7414, were performed using LC-MS/MS. Interestingly for all three species, it is the same PsaL protein, encoded by *psaL* in *psaF/J/L*, that was observed in both the tetrameric and trimeric PSI of these three strains (Fig. S5). Moreover, genome sequencing of TS-821 revealed only a single form of *psaL*. Although PCC 7414 encodes two copies of the *psaL* gene (Fig 2a), only the PsaL encoded by *psaL* in the *psaF/J/L* structure was found in either PSI form, isolated under our experimental conditions. This proteomic result reinforced the possibility that the *psaL* encoded in *psaL/I* in PCC 7414 is an example of far-red light responsive PSI genes (Gan et al., 2014; Gan and Bryant, 2015). These far-red light responsive genes are organized differently in the genome. Moreover, when we build a maximum-likelihood tree of PsaL, the group of far-red light responsive forms of PsaL form a distinct clade (in red text) as shown in Fig. S4b. Phylogenetic analysis revealed the PsaL within the *psaF/J/L* loci of the HCR forms a single clade. However, the observation that the same PsaL is observed in both tetrameric and trimeric PSI in TS-821, PCC 7428, and PCC 7414 suggests that the unique PsaL encoded in *psaF/J/L* is necessary, yet not sufficient, to direct PSI tetramer formation.

### PsaL Replacements affect PSI Oligomerization

To investigate precisely what structural property of PsaL is key to the determination of the PSI oligomeric state, we performed gene replacement experiments in the mesophilic cyanobacteria *Synechocystis* sp. PCC 6803. Replacing WT PsaL in *Synechocystis* sp. PCC 6803 by homologous recombination with either the TS-821 PsaL or *Arabidopsis* PsaL resulted in monomeric PSI in both mutants (Fig. 3a, b). When western blotting was performed on these monomeric forms of PSI, it was clear that they too contained PsaL bound to PSI (Fig. 3c, d). This observation suggests that the monomerization of PSI in these transgenic lines is not due to the lack of PsaL expression and/or assembly. These findings suggest that subtle changes in the PsaL structure alone can lead to the monomerization of trimeric PSI in cyanobacteria, as found with the predominant monomeric PSI in PCC 6605 (Figs 1c, S3a).

**Fig. 3.**
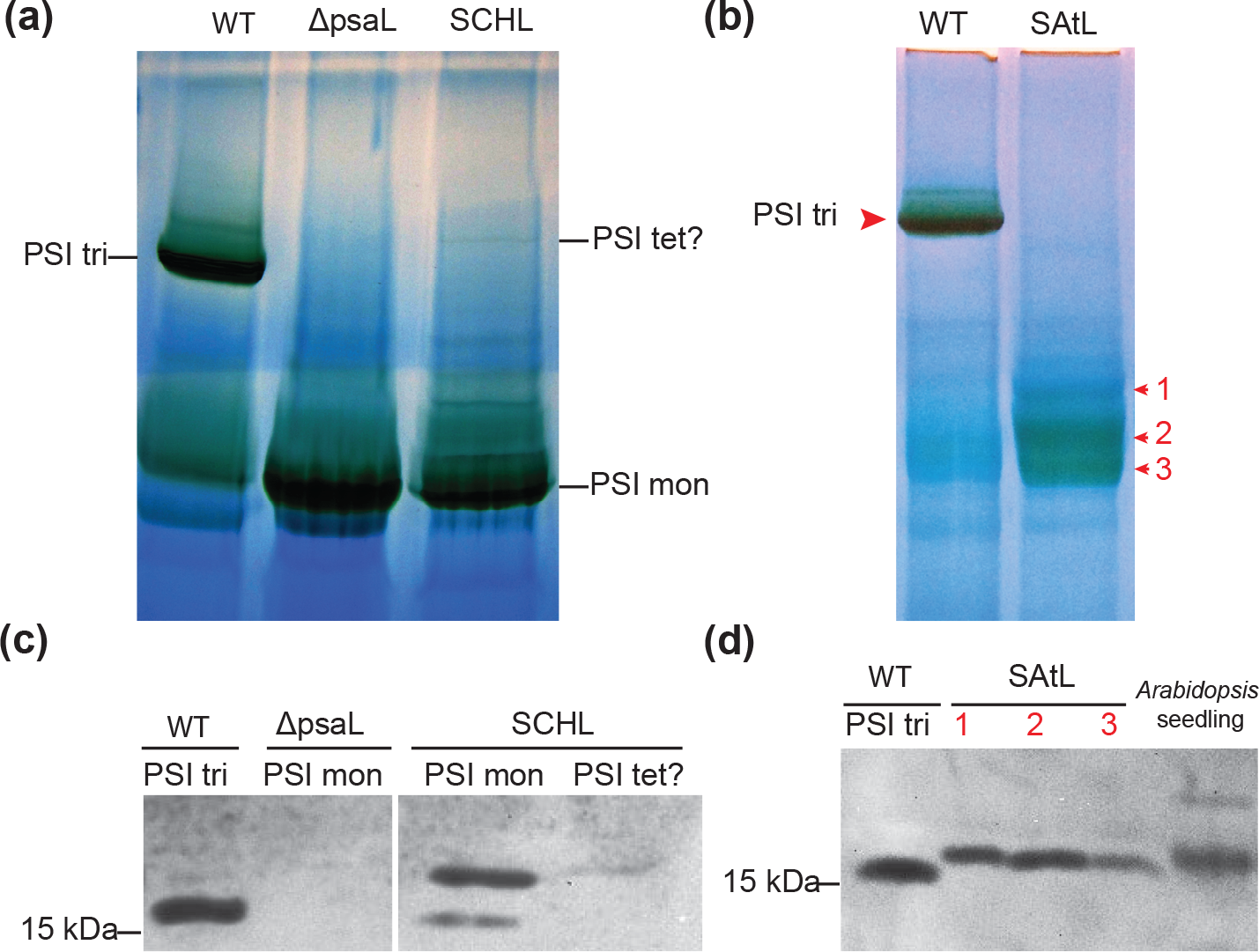
BN-PAGE and western blot analyses of PSI oligomers in *Synechocystis* sp. PCC 6803 expressing different PsaL. **(a)** BN-PAGE of solubilized thylakoid membranes from *Synechocystis sp*. PCC 6803 WT, *∆psaL*, and expressing TS-821 PsaL (SCHL). PSI samples were sliced out for from the gel for following SDS-PAGE and western blot to detect the presence of PsaL. Each lane was loaded with solubilized membrane containing 16 μg Chl. **(b)** BN-PAGE of solubilized thylakoid membranes from *Synechocystis sp*. PCC 6803 WT and mutant expressing *Arabidopsis thaliana* PsaL (SAtL). **(c)** Western blot of the PSI samples from BN-PAGE gel **a**. PSI trimer (tri) from WT, PSI monomers (mon) from *∆psaL* and SCHL, as well as potential PSI tetramer (tet?) were analyzed. **(d)** Western blot of the PSI bands isolated from BN-PAGE gel **b**. *Arabidopsis thaliana* seedling whole plant was used as a control.

Our earlier work on TS-821 speculated that the loop sequence of PsaL between the second and third transmembrane helices might be a key part in PSI oligomer and PsaL evolution (Li et al., 2014). While this loop sequence structure was not resolved in the trimeric PSI crystal structure from *T. elongatus* (PDB 1JB0), later improved plant PSI monomer structures showed the interaction between this loop sequence with PsaH (Qin et al., 2015, Mazor et al., 2015). Noticing that TS-821 PsaL has an unusual multi-proline motif in this loop sequence, we examined the conservation of this loop sequence between PsaL second and third transmembrane helices for all HCR we studied for their PSI oligomeric states (Fig. S6). By building a LOGO plot of these regions of PsaL from all studied 38 of the HCR, we identified a conserved proline-rich motif, often as NPPxP followed by PNPP (Fig. S6). Many of the trimeric PSI containing sequences lack this motif. However, interestingly this motif is shared by both copies of PsaL in PCC 6605 (Fig. S6) which has predominantly monomeric PSI (Figs 1c, S3a). These observations suggest a similar mechanism of trimeric PSI destabilization in PCC 6605 involving a similar multi-proline motif as observed in this part of PsaL in the heterocyst-forming cyanobacteria.

### Influence of Environmental Factors

It is now clear that the formation of PSI tetramers is restricted to a specific subset of cyanobacteria. Despite our ability to “map” this trait to the HCR, there was no clear understanding as to the physiological or evolutionary role that drives this change in oligomerization. Since most heterocyst-forming cyanobacteria have tetrameric PSI, the impact of nitrogen sources and presence of heterocysts on PSI tetramer formation was investigated. We grew three heterocyst-forming strains (PCCC 7414, PCC 7120, and PCC 7122) in both nitrate depleted BG-11 media (BG-11_0_) and BG-11 supplemented with NH_4_^+^. Based on the similar PSI pattern observed with BN-PAGE (Fig. S7a) as observed in those cultured in standard BG-11 media (Fig. S2j, k), it indicates that the nitrogen sources did not affect the presence of tetrameric PSI. Similar result was also observed for *Fischerella* sp. UTEX LB 1829 (BG-11_0_ vs BG-11, Fig. S2e). These strains did not develop heterocysts with supplemented NH_4_^+^, indicating the presence of tetrameric PSI in vegetative cells. The presence of tetrameric PSI in unicellular strains such as TS-821, PCC 7509 and PCC 7428 (Figs 1a, S2a, b) also agrees with a PSI tetramer independent from heterocyst development, even though tetrameric PSI was shown in heterocysts (Cardona et al., 2009). In the case of *Nodularia* PCC 73104, the small amount of PSI tetramer was only observed in BG-11 medium (Fig. S2j), while the addition of NH_4_^+^ or salt in the culture media appeared to diminish the tetrameric PSI (Fig. S7c). The special case of *Nodularia* PCC 73104 needs more studies to elucidate the significance of such response, since *Nodularia* are usually found in brackish or oceanic environments with some strains do not abolish their heterocysts in the presence of NH_4_^+^ (Vintila and El-Shehawy, 2007; Vintila and El-Shehawy, 2010).

In addition to nitrogen source, growth temperature had little, if any, effect on the presence of tetrameric or trimeric PSI (Fig. S7b) on the four different cyanobacteria investigated. For the impact of salinity, two strains of *Nodularia*, heterocyst-forming cyanobacteria isolated from brackish or saline water, were cultured in both fresh water and artificial seawater media (ASNIII) we observed little change in the amount of the tetramer/dimer (Fig. S7c). At the same time, salinity did not affect the presence and amount trimeric PSI in *Pseudanabaena* sp. B SP48 (Fig. S3e). Together these results suggest that none of the factors presented above, i.e, nitrogen source, heterocyst development, temperature stress, and salinity, has any effect on the formation of tetrameric and trimeric PSI.

### PSI Tetramer in Response to Light Intensity

In contrast to other environmental factors, upon altering the light intensity during growth we saw a clear effect on the ratio of the PSI oligomeric forms. Our study on TS-821 showed that in response to light intensity change from low light (LL) to high light (HL), the tetrameric PSI accumulated, while the PSI trimer decreased (Semchonok et al., 2016). To confirm that light intensity was a common factor determining PSI tetramer/trimer ratio, we investigated the HL response of additional cyanobacterial strains. We observed an increase in quantity and stability of the tetrameric PSI for both PCC 7428 and PCC 7414 under HL condition (Fig. 4). In addition, it was clear that the amount of trimeric PSI was much higher for PCC 7428 when grown at low light (LL) levels (Fig. 4a, b). The difference observed in PCC 7414 under HL versus LL was slightly less than that observed in PCC 7428 (Fig. 4c, d) possibly due to self-shading within the intertwined filaments in PCC 7414 (Fig. S8). Yet these results point to a common HL response, accumulating tetrameric PSI and suppressing trimeric PSI in strains having both forms of the PSI oligomers. Therefore, it is reasonable to speculate that tetrameric PSI is better adapted to HL than trimeric PSI.

**Fig. 4.**
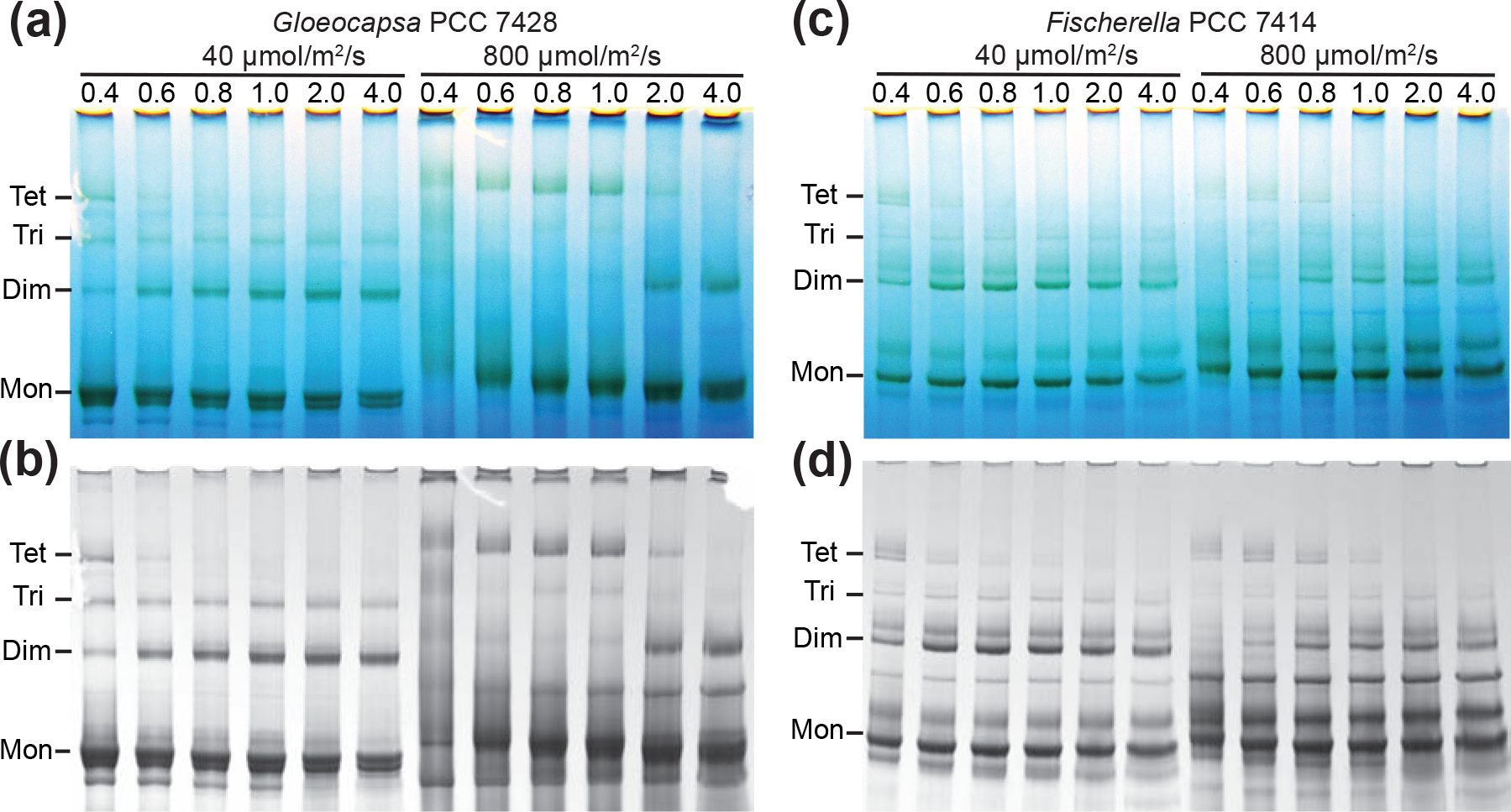
BN-PAGE analyses of cyanobacterial PSI oligomeric states under different light intensities. **(a)** and **(c)** show the unstained BN-PAGE analyses of solubilized thylakoid membranes from PCC 7428 and PCC 7414 respectively. The left lanes are the low light (40 μmol/m^2^/s) and the right lanes are the high light (800 μmol/m^2^/s). **(c)** and **(d)** present the same BN-PAGE gels as **a** and **c** respectively after staining with Coomassie to enhance clarity. Thylakoid membranes with 0.2 mg/mL Chl were solubilized in different concentrations of DDM (w/v, %), labeled on top of each lane. ~ 1.6 μg Chl were loaded for each condition. Main bands corresponding to PSI monomer (Mon), Dimer (Dim), Trimer (Tri), and Tetramer (Tet) are labeled.

We have performed a more detailed study of PSI oligomer profiles for TS-821 cultured under several different light intensities, which supported the hypothesis that tetrameric PSI is an adaption to HL. As shown in Fig. 5, the stability and relative quantity of the PSI tetramer in TS-821 increased as the intensity of light is raised from 50-800 mol/m^2^/s. (Fig. 5a-g). When the relative quantities of PSI tetramer (percentage over all PSI oligomers) was plotted versus light intensities, it was clear that the apparent maximum increased with light intensity (0.4% DDM, Fig. 5g). The stability of PSI tetramer, indicated by the relative quantity at higher detergent concentration (0.8% DDM, Fig. 5g), also was enhanced by higher light intensities. In contrast, the relative amount of trimeric PSI decreases nearly linearly as the light intensity increases (Fig. 5h). This ability to alter the oligomeric form of PSI is not found in all species of *Chroococcidiopsis*, as indicated by the observation that the absence of PSI tetramer in PCC 7203 was not affected by light intensity (Fig. S7d).

**Fig. 5.**
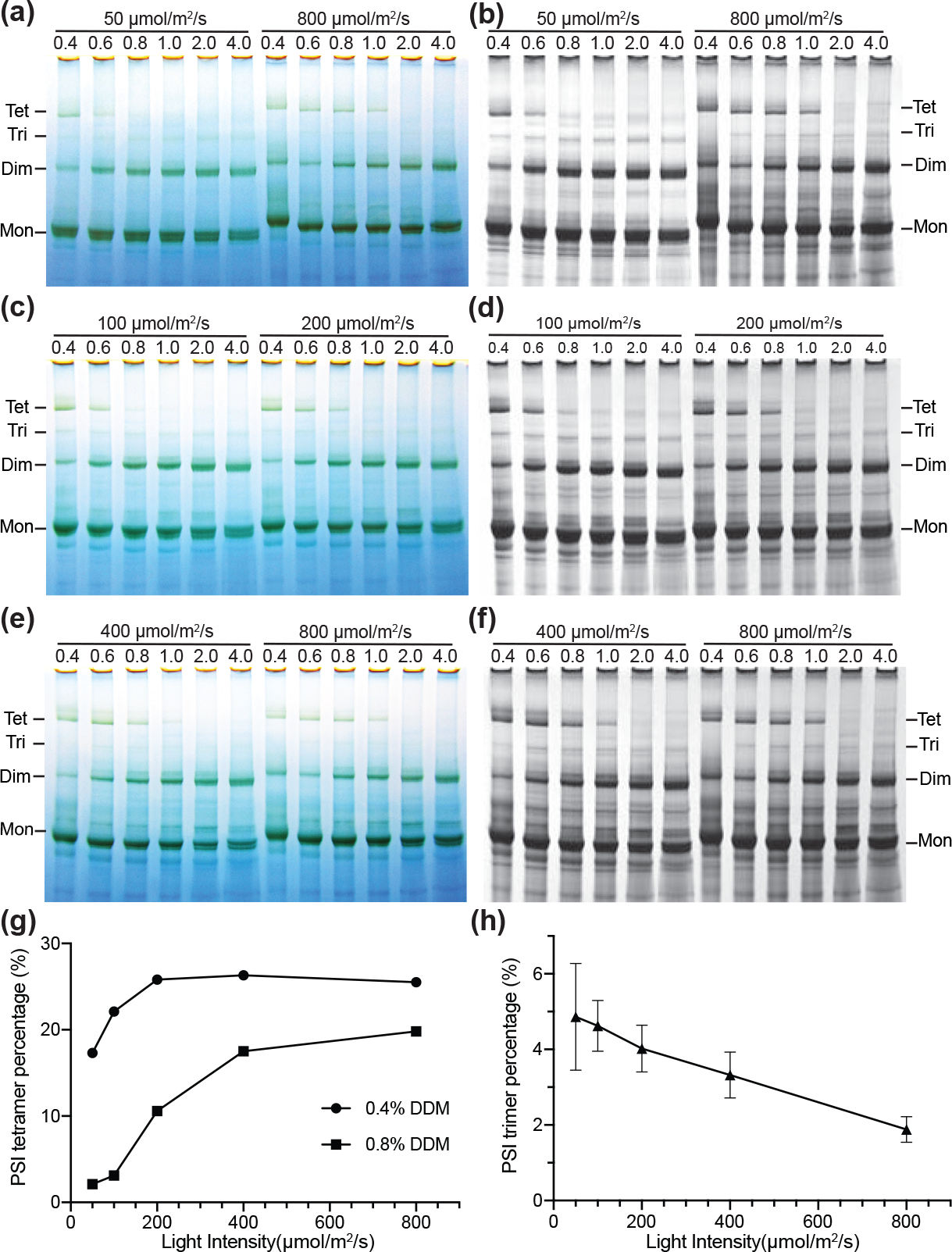
Comparison of TS-821 PSI oligomeric profiles under different light intensities. **(a) (c)** and **(e)** BN-PAGE of solubilized thylakoid membrane from TS-821 cultured under different light intensities. Coomassie stained images are shown as in **(b) (d)** and **(f)** for clarity and quantification. Thylakoid membranes with 0.2 mg/mL Chl were solubilized in different concentrations of DDM (w/v, %), labeled on top of each lane. ~ 1.6 μg Chl were loaded for each condition. Main bands corresponding to PSI monomer (Mon), dimer (Dim), trimer (Tri), and tetramer (Tet) are labeled. **(g)**, Plots of relative quantities of PSI tetramer under different light intensities. Two solubilization conditions, 0.4% and 0.8% DDM, are presented to show the observed maximum quantities and the stabilities of PSI tetramer under different light intensities. **(h)**, Plot of relative quantities of PSI trimer under different light intensities. Error bar: standard deviation (n=5 solubilization conditions).

To investigate whether tetrameric PSI in TS-821 undergoes a shift in pigment composition or biochemical/biophysical changes under different light intensities, absorption spectra and 77K fluorescence spectra were compared for different PSI oligomers isolated from cells grown under different light conditions (Figs 6a, b, S9). While the 77K emission spectra of these isolated PSI oligomers did not show significant differences (Fig. S9), it is clear that the PSI tetramer under HL has an additional absorption in the range around 440-560 nm, compared with tetrameric PSI under LL (Fig. 6a, b). The difference spectrum from these two samples showed that at least three discernable peaks (450, 490, and 520 nm) (Fig. 6b). The absorption spectrum of the LL PSI tetramer also showed a similar additional absorption around 400-500 nm when compared with the PSI trimer under LL (Fig. S9).

**Fig. 6.**
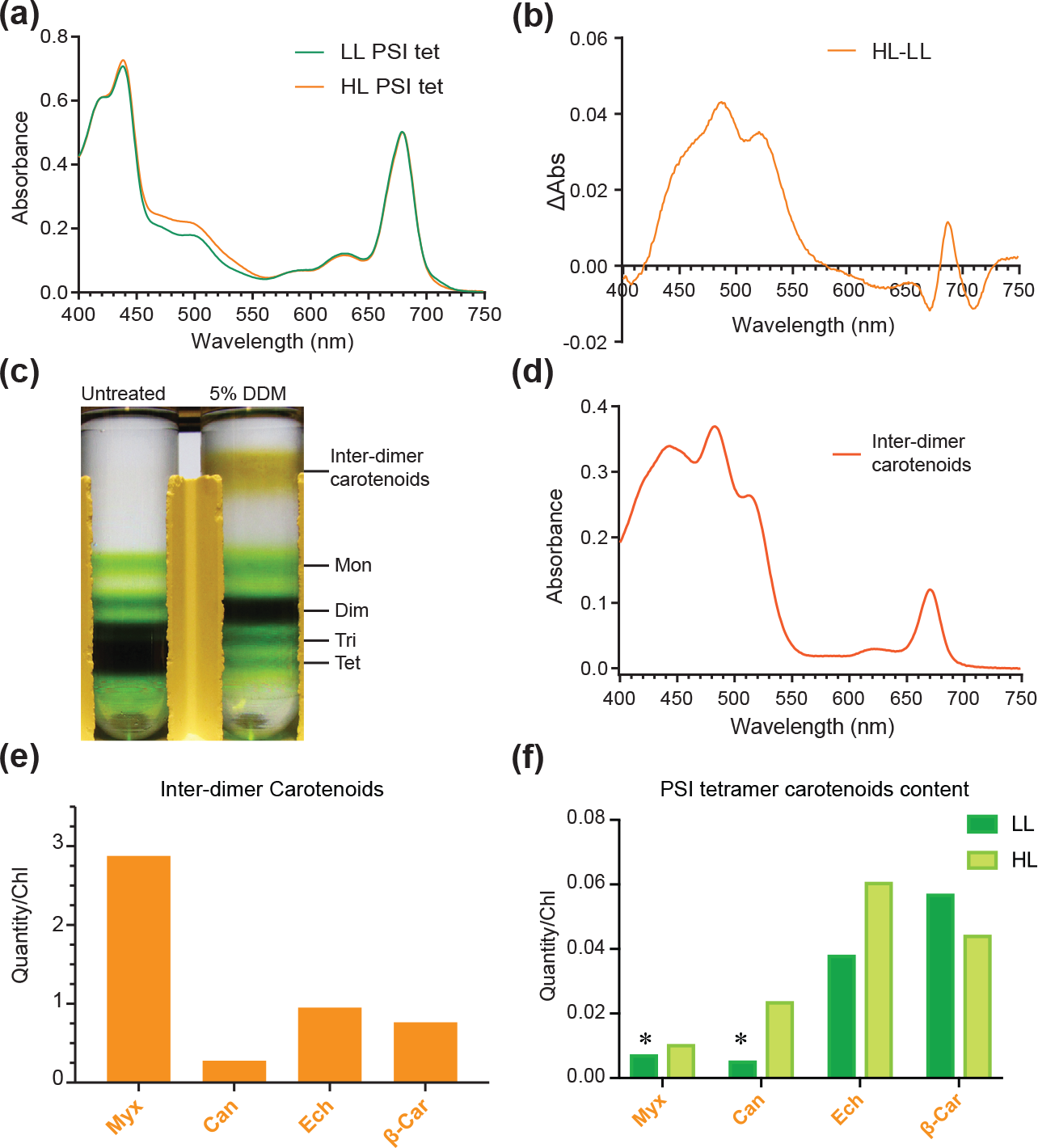
The impact of light intensities on tetrameric PSI in TS-821. **(a)** Absorption spectra of PSI tetramer isolated from TS-821 grown under low light (LL) and high light (HL). **(b)** Absorption spectrum difference between HL and LL PSI tetramer calculated from spectra **a**. **(c)** Sucrose density gradient ultracentrifugation of isolated PSI tetramer. The sample treated with 5% DDM is compared with untreated PSI tetramer. Main bands corresponding to inter-dimer carotenoids, PSI monomer (Mon), Dimer (Dim), Trimer (Tri), and Tetramer (Tet) are labeled. **(d)** Absorption spectrum of inter-dimer carotenoids isolated from treated PSI tetramer after sucrose density gradient ultracentrifugation **c**. **(e)** Quantitative analysis of carotenoids released upon inter-dimer carotenoids. **(f)** Quantitative comparison of carotenoids in PSI tetramer isolated under high light (HL) and low light (LL) conditions. The ratios of carotenoids to Chl are presented. The detected carotenoids include myxoxanthophyll (Myx), canthaxanthin (Can), echinenone (Ech) and β-carotene (β-Car). Asterisks (*) denote the detected quantities are below limit of quantification.

Interestingly, when the PSI tetramer is treated with 5% DDM and dissociated into dimers, a very prominent amount of carotenoids stayed on the top layer of the sucrose gradient after separation by centrifugation (Fig. 6c). It is very likely that these carotenoids are associated with the inter-dimer interface and may even reside in the center of the tetrameric structure, only being released upon dissociation. Interestingly, the absorption spectrum of released carotenoids (Fig. 6d) resembled the increased absorbance observed in the HL tetramer (Fig. 6b). Moreover, spectral analysis of the tetrameric PSI in both TS-821 and PCC 7414 reveal an increase in absorption in the range of 400 ~550 nm when compared with the PSI dimer and trimer (Fig. S10c, d), and is quite similar to the inter-dimer carotenoids absorption spectrum (Fig. 6d). These results suggest that tetrameric forms of PSI are generally rich in carotenoids, compared to trimeric and dimeric forms of PSI. When these inter-dimer carotenoids were isolated and analyzed by HPLC, we identified novel PSI cofactors: myxoxanthophyll, canthaxanthin, echinenone, in addition to known cofactors: chlorophyll (Chl) and β-carotene (Figs. 6e, S11a). The semi-quantitative comparison between the HL PSI tetramer and LL PSI tetramer also revealed that tetrameric PSI harbored more myxoxanthophyll, canthaxanthin, echinenone and less β-carotene (per Chl) under HL (Figs. 6f, S11).

## Discussion

Our results indicate that tetrameric PSI is a characteristic shared by most, if not all, heterocyst-forming cyanobacteria and their closest unicellular relatives. The distribution of tetrameric PSI in this clade of cyanobacteria points out a shared physiological characteristic between the two morphologically distinct cyanobacteria. The prior classification of PCC 7428 and TS-821 would place these cyanobacteria in Section II based on their reproduction by multiple fission giving rise to baeocytes (Rippka et al., 1979), but our current work on these PSI-dimer/tetramer containing unicellular cyanobacteria would support a different placement. Phylogenetic analyses based on 16S rRNA gene as well as a more universal set of conserved genes suggest that these tetramer-forming cyanobacteria are closely related to heterocyst-forming cyanobacteria (Fewer et al., 2002; Shih et al., 2013). However, no further physiological or biochemical evidence had been reported to support the close evolutionary linkage between the two groups. In this study, we showed similar physiological and biochemical properties of the PSI oligomers observed in those two groups, providing further support of a phylogenetic placement of the PSI-dimer/tetramer unicellular cyanobacteria close to heterocyst-forming strains, as one clade, HCR.

Additionally, the phylogenetic analysis provided a clear classification of PsaL in correlation to PSI oligomeric state (Fig. S4). With given cyanobacterial *psaL* sequences, the major PSI oligomeric states can be predicted, even though specific strains may vary from the major correlation between PsaL and PSI oligomer, such as PSI monomer found in heterocyst-forming cyanobacteria (Fig. S1). The formation of the tetrameric structure may be a result of changes in the PsaL subunit, as proposed in earlier work (Li et al., 2014), and it genomic location in relation to *PsaF/J* which can be an element for expression regulation and PSI assembly control. Yet changes in PsaL are insufficient for PSI tetramer formation, consistent with the observations of PSI monomer in PCC 6803 expressing TS-821 PsaL. Besides differences in PsaL, other factors such as carotenoids and PSI assembly chaperones, may contribute to the formation of tetrameric PSI.

The discovery of a pool of PSI inter-dimer carotenoids also suggests the physiological significance of tetrameric PSI over trimeric PSI, since these additional carotenoids may protect PSI from excessive influx of photons under HL. This inference is further supported by the fact that the carotenoid content, as well as the relative quantity and stability of PSI tetramer increased under HL (Figs 5, 6). Even under LL, PSI tetramer has higher content of carotenoids compared with PSI trimer (Fig. S10). This is in line with the observations of antennae cooperativity between the three *T. elongatus* PSI monomers to yield an increased optical cross-section of the trimer (Iwuchukwu et al., 2010; Baker et al., 2014), while tetrameric and dimeric PSI did not show significant cooperativity among the monomers (Li et al., 2014). Altogether, tetrameric PSI by itself, compared to trimeric PSI, is a less efficient form of PSI in light harvesting but a better form of PSI in terms of dissipating excessive photonic energy under HL conditions, thereby avoiding photoinhibition. Thus, these cyanobacteria can alter the oligomeric state of PSI as light increases, while also increasing the content of the photoprotective carotenoids. It is not clear if it is the increase in the carotenoids that facilitate the formation of the tetramer or vice versa, yet it is attractive to speculate that the tetrameric structure provides a mechanism to accumulate carotenoids possibly via an unknown and/or novel carotenoid binding mechanism. The light responsive adaptive behavior of the PSI oligomeric state and the ability of a PSI tetramer adjusting its carotenoid content, represents an early evolutionary and unique photo-protective mechanism in HCR.

Our results support the hypothesis that PSI oligomers evolved from trimeric to tetrameric in cyanobacteria *en route* to exclusively monomeric in plants and green algae. By comparing PSI trimeric structure from cyanobacteria and PSI monomeric structure from plants, it was proposed that PSI evolved from a PSI trimer to PSI monomer due to the presence of PsaH, preventing the formation of PSI trimer (Amunts and Nelson, 2009). However, the presence of exclusively monomeric PSI in cyanobacteria (Fig. S2), red algae and heterokonts (Alboresi et al., 2017) without the involvement of PsaH suggests that PSI monomerization preceded the introduction of the PsaH subunit. Changes in PsaL appeared sufficient for PSI monomerization notably with *Synechocystis* expressing *Arabidopsis* PsaL exhibiting exclusively monomeric PSI (Fig. 3b). Therefore, the plant PsaL appeared more closely related to cyanobacterial PsaL in tetrameric/monomeric PSI. The presence of exclusively monomeric PSI in heterocyst-forming cyanobacteria (Fig. S2) or mostly monomeric PSI in examples such *Chroococcidiopsis* sp. PCC 7203 (Figs 1a, S2) suggests that the monomerization of PSI in plants and green algae shared the same or closely related route to that of PSI tetramer formation over the course of evolution.

Exposure to HL may be the driving force for a change in PSI oligomerization. The shift of PSI oligomeric structure from trimeric to tetrameric under HL intensities supports that the HL condition associated with terrestrial environments may have been the selection pressure for PSI tetramer formation. It had been observed that *Synechocystis* sp. PCC 6803, although having mainly trimeric PSI, accumulates more PSI monomer when grown under HL (Wang et al., 2008), which suggests that monomeric PSI may also be advantageous over trimeric PSI under this increased light intensity. Our study on *Chroococcidiopsis* sp. PCC 7203, which had mostly monomeric PSI and some dimeric PSI, showed no obvious oligomeric shift when transferred from LL to HL (Fig. S7). Whether monomeric PSI is a further adaptation to HL remains to be investigated.

Nevertheless, it is reasonable to speculate that early plastid ancestors needed to be able to cope with HL irradiance as they moved from relatively light limiting marine/aquatic settings onto the surface of a terrestrial environment. Compared with the PSI trimer, the PSI tetramer or monomer seem to be more adapted to such light intensity. These PSI oligomers are also well suited for the addition of other PSI subunits such as PsaH and light harvesting complexes that bind to the inter-monomer faces of PSI. Finally, the observation of a very similar PSI tetramer in the glaucophyte *Cyanophora paradoxa* (Watanabe et al., 2011) suggests that the closest cyanobacterial relative to plastid ancestors may have also had tetrameric PSI, reinforcing the gene homologies and phylogenomic conclusions that heterocyst-forming cyanobacteria are the most closely related to the plastid ancestor (Dagan et al., 2013).

## Materials and Methods

### Cyanobacterial growth conditions

The regular growth conditions for three control strains*, Synechocystis* sp. PCC 6803 (WT and mutants), *Chroococcidiopsis* sp. TS-821 (TS-821) and *T. elongatus* BP-1 were described in earlier study (Li et al., 2014). Cyanobacteria from UTEX were maintained as recommended by the culture collection center at University of Texas, Austin, USA, and the Pasteur Culture Collection of Cyanobacteria at the Institut Pasteur, Paris, France (Dataset S1). To achieve sufficient cell mass for PSI oligomer identification, the cyanobacteria were cultured in feasible media with light intensity in the range of 20~100 μmol photons /m^2^/s (Dataset S1). The modified ASN III medium contains BG-11 medium with addition of ASNIII media major salts (25 g/L NaCl, 3.5g/L MgSO_4_ × 7H_2_O, 2.0g/L MgCl_2_ × 6H_2_O, 0.5 g/L KCl).

To study the physiological significance of PSI tetramer in cyanobacteria, non-standard culture conditions were tested. To investigate the effect of nitrogen sources, some heterocyst-forming cyanobacteria were cultured separately in BG-11, BG-11_0_ or BG-11+NH_4_^+^ (BG-11 supplemented with 1 mM NH_4_Cl two days before cell harvesting). To test the effect of temperature, some thermophilic cyanobacteria were cultured at 45 °C versus 37 °C. In addition, a culture of *Fischerella muscicola* PCC 7414 grown at 37 °C was incubated at 24 °C before harvesting. To investigate the effect of salinity, some marine cyanobacteria were cultured in both BG-11 and modified ASNIII media.

The standard high light (HL) and low light (LL) comparison experiments for TS-821, PCC 7414, PCC 7428, and PCC 7203 were done as described before (Semchonok et al., 2016). To eliminate the potential stress from cell culture density and/or difference from lighting sources, a systematic analysis of how TS-821 respond to different level of light intensity was done by culturing similar inoculum in photobioreactor for two days under five light conditions (50, 100, 200, 400 and 800 μmol photons/m^2^/s). For each light condition, half of the light intensity was provided by white LED light and the other half was from red LED. The mid-log phase TS-821 cell cultures (OD_680_ 0.6-0.8) under white light fluorescent light were inoculated to the 25 L bioreactor to starting OD_680_ at 0.04 (measured by equipped fluorometer probe from Photon Systems Instruments, Drazov, Czech Republic) before turning on the bioreactor lights. Cells were harvested and kept in −20 °C before PSI oligomer analysis.

### Thylakoid membrane isolation

For thylakoid membrane preparation, cells are lysed by French Press. Cyanobacterial cell pellets were washed in buffer A (50 mM MES-NaOH, pH 6.5, 5 mM CaCl_2_, 10 mM MgCl_2_) and then pelleted by centrifugation (Watanabe et al., 2009). Cell pellets were resuspended in lysis buffer (buffer A containing 0.5 M sorbitol) and homogenized before French Press. In the case of *Scytonema crispum* UTEX LB 1556, cell aggregates were broken into small fragments by grinding in liquid nitrogen before resuspension in lysis buffer and homogenization. Homogenized cell suspensions were processed through the French Press three times at 1500 or 2000 psi (10 to 14 MPa), then unbroken cells were removed by pelleting at 10,000 g for 5 min.

In most cases, thylakoid membrane pelleting from cell lysate was done by centrifugation at 40,000 rpm (193,000 g Type 50.2 Ti, Beckman) for 30 min at 4 °C. Thylakoid membrane pellets were resuspended in buffer A + 12.5% glycerol and homogenized before storing at −20 °C or −80 °C. For HL versus LL experiments, thylakoids were washed in buffer A once and pelleted at 193,000 g for 15 min before final resuspension. In some special cases where cell mass is low and large membrane fragments are ready to be pelleted at lower g-force, thylakoids were resuspended after spinning at 10,000 g for 5 min. Chl concentration was determined as described previously (Iwamura et al., 1970).

### PAGE analyses and Western blot

In most cases, 4-16% BN-PAGE gels (Invitrogen) were used to analyze solubilized thylakoids or isolated photosystems according to the user manual and references (Schägger and von Jagow, 1991; Wittig et al., 2006). In addition, some of the BN-PAGE gels were homemade to analyze samples with large loading amount. To analyze PSI oligomeric states, thylakoid membranes were solubilized in different concentration of detergent n-dodecyl-β-maltoside (DDM) (Glycon, Germany) at 25°C for 1.5 h. Insoluble material was removed by centrifuging 180,000 g for 5 min or 98, 000 g for 10 min at 4 °C. Supernatants were taken out for BN-PAGE analysis. To identify the photosystems after BN-PAGE, a second dimension Tris-Tricine SDS-PAGE was used as described previously (Li et al., 2014).

To detect the presence of PsaL in *Synechocystis* mutants, Western blot was done after SDS-PAGE. Protein bands were transferred from polyacrylamide gels to a 0.45 μm PVDF membrane (Millipore) using blotting cassette (idEA, Minneapolis, MN). Membranes were blocked using TBS-T buffer + 3% NFM (non-fat milk powder) at room temperature for 1 hour. Primary antibody antisera were diluted from 1: 5000 in TBS-T + 3% NFM depending on the efficiency of antisera before treating the blocked membranes overnight at 4 °C. Secondary antibody (goat anti-rabbit) conjugated to HRP was diluted in TBS-T + 3% NFM down to 1:50,000 before treating the washed membranes for 1 hour at room temperature. The targeted proteins were then detected using chemiluminsence HRP substrate (Millipore) with signal recorded using ChemiDoc XDS system (Bio-Rad).

### Antigen design and antibody production

PsaL sequences are not well conserved for antigen design from a single consensus sequence. To achieve one antigen for all PsaL of interest in this study, i.e. *<u>C</u>hroococcidiopsis* sp. TS-821, *<u>A</u>rabidopsis*, *<u>T</u>. elongatus* BP-1, and *<u>S</u>ynechocystis* sp. PCC 6803, a fused protein, CATSPsaL containing fragments of different PsaL was designed. To predict the epitopes, Kolaskar & Tongaonkar Antigenicity’s methods (Kolaskar and Tongaonkar, 1990) on IEDB Analysis Resource (http://tools.immuneepitope.org/tools/bcell/iedb_input) and ABCpred (Saha and Raghava, 2006) (http://www.imtech.res.in/raghava/abcpred/index.html) were used. The antigen sequence is aligned against different PsaL fragments and its predicted antigen to ensure the high probability of getting antibody for each PsaL (Fig. S1). In addition to the N-terminal fragments, the loop insertion between second and third trans-membrane helices of TS-821 PsaL was also used to achieve high antigenicity for TS-821.

Synthetic *CATSpsaL* gene (IDT, San Jose, CA) was cloned into pTYB2 (NEB) plasmid. The expression plasmid DNA was then transformed into *E. coli* ER2566 for antigen expression. IMPACT™ Chitin Resin (NEB) was used as column matrix for antigen purification. Antibody was produced at Pocono Rabbit Farm (Canadensis, PA) using the 91-day protocol on two rabbits. For PsaL antibody production, the purified 1 mg/mL antigen was used as inoculum.

### PSI isolation and characterization

Sucrose Density Gradient Centrifugation (SDGC) was used for PSI isolation as described previously (Li et al., 2014). SDGC was also used to study PSI oligomeric states, as a parallel or complementary experiment to BN-PAGE profiling. Thylakoid membranes containing 0.4 mg/mL Chl were solubilized in 0.6~1% DDM. After removing insoluble fragments, the solubilized membranes containing 1 mg Chl were loaded on 10-30% sucrose density gradient in buffer A containing 0.01% DDM. This method showed comparable result with BN-PAGE analysis. For large loading of solubilized membrane (containing 1~3 mg Chl) on a sucrose density gradient, centrifugation was done twice with the first 20~24-hour spin followed by dialysis and another 24-hour spin (SW32Ti, 30,000 rpm). Lower ionic strength such as 0.4 × buffer A in sucrose density gradient for PCC 7414 PSI oligomers was tried and showed no observable difference for PSI profiles. Inter-dimer carotenoids were isolated after SDGC following treatment of PSI tetramer with 5% DDM for at least one hour at room temperature. Isolated PSI samples were dialyzed before analysis. For the proteomic comparison between PSI trimers and PSI tetramers, isolated PSI oligomers were further purified by BN-PAGE and sliced out for analyses.

Low temperature fluorescence spectra were acquired as described before (Li et al., 2014). Absorption spectra of isolated proteins and pigments were taken using Cary 300 UV-Vis spectrometer (Agilent). The sample buffers were used as blanks. For PSI oligomer absorption spectra comparisons, A_680_ was adjusted close to 0.5 or 1. The PSI absorption differences were calculated after normalizing the data at A_680_. Electron microscopy and single particle analyses were done as described in previous studies (Li et al., 2014; Watanabe et al., 2014). Pigment analyses were done using HPLC by DHI (Denmark) (Van Heukelem L and Thomas CS, 2001).

### *psaL* gene cloning and TS-821 genome sequencing

Most cyanobacterial DNA extracted for *psaL* cloning were done as described in previous study (Li et al., 2014). TS-821 DNA for genome sequencing as well as some cyanobacterial DNA for *psaL* cloning was extracted using the NucleoBond^®^ DNA isolation kit. For *psaL* cloning of heterocyst-forming cyanobacteria with unknown genome data, different primer sets (Tables S1, S2) from *psaF* and *gmk* genes were used to amplify the *psaL* gene and its flanking region. The cyanobacteria of interests were classified using their known DNA sequences or genus names to facilitate picking primer pairs. PCR products were ligated into pJET vectors using CloneJET™ PCR cloning kit. Plasmids containing cloned *psaL* were sequenced using Sanger sequencing at UT Genomics Core (Knoxville, TN).

TS-821 culture was purified as described in earlier study (Rippka et al., 1979) before DNA extraction for genome sequencing. The DNA was submitted to UT Genomic Core (Knoxville, TN) for sequencing using Illumina^®^ MiSeq Kit V2 with 2 × 250 bp read length. The resulted FASTQ files were analyzed using SPAdes (version 3.7.1) (Bankevich et al., 2012). The assembled contigs were submitted to PATRIC (Wattam et al., 2014) for annotation, which uses the RAST (Overbeek et al., 2014) system.

### Phylogenetic and *psaL* analyses

The species tree was generated by a concatenation of twenty-nine conserved proteins selected from the phylogenetic markers previously validated for cyanobacteria using a Maximum Likelihood method as described previously (Calteau et al., 2014). Protein sequences were aligned using MAFFT v7.307 (Katoh and Standley, 2013), then ambiguous and saturated regions were removed with BMGE v1.12 (with the gap rate parameter set to 0.5) (Criscuolo and Gribaldo, 2010). The best fitting model of amino acid substitution for this dataset was selected with ProtTest v3.2 (Darriba et al., 2011). A Maximum-Likelihood phylogenetic tree was generated with the alignment using PhyML 3.1.0.2 (Guindon et al., 2010) using the LG amino acid substitution model with gamma-distributed rate variation (six categories), estimation of the proportion of invariable sites and exploring tree topologies. 100 bootstrap replicates were performed. The phylogenetic trees were displayed and annotated using the interactive tree of life (iTOL) online tool (Letunic and Bork, 2016).

Genomic context of *psaL/I/F/J* genes and PsaL protein sequences were extracted from the MicroScope platform (Vallenet et al., 2017), public databases, and data generated in previous study (Schirrmeister et al., 2015) or this study (Dataset S1). The LOGO plot for the PsaL linker was generated using WebLogo 3 website (http://weblogo.threeplusone.com) (Crooks et al., 2004).

### Construction of *Synechocystis psaL* mutants

Some *psaL* mutants were obtained in previous study (Li et al., 2014) with low expression efficiency. To improve the expression level of exogenous PsaL, the codon optimized synthetic *psaL* DNAs in pBSK vector were obtained from IDT (San Jose, CA). Similar methods were used for cloning and *Synechocystis* transformation as described earlier (Li et al., 2014). The mutants are named as *Synechocystis* sp. PCC 6803 expressing <u>S</u>ynthetic *<u>Ch</u>roococcidiopsis* sp. TS-821*/<u>A</u>rabidopsis <u>t</u>haliana psa<u>L</u>* (SCHL/SAtL).

## Acknowledgments

Support to B.D.B., M.L., and J.T.N. has been provided from the Gibson Family Foundation, the Bredesen Center for Interdisciplinary Research and Education, the Tennessee Plant Research Center, a UTK Professional Development Award, the Dr. Donald L. Akers Faculty Enrichment Fellowship to B.D.B. and National Science Foundation support to B.D.B. (DGE-0801470 and EPS-1004083). M.L. has been supported as a CIRE Fellow at University of Tennessee, Knoxville. A Professional Development Award from the Graduate School at UTK supported travel of B.D.B. to the Netherlands and to the Institut Pasteur. NWO Chemical Sciences supported work at University of Groningen. J.P.W. has been supported from NIH P30 DK063491. The Institut Pasteur supported Pasteur Culture Collection of Cyanobacteria. We are also grateful to Y.I. Park for the use of cyanobacterial genome of PCC 7124. We would like to thank Mr. Nathan G. Brady for helpful comments on the manuscript.

## Author contributions

M.L., M.G. and B.D.B designed the research. M.L. carried out most of the biochemistry and molecular biology experiments. A.C did the phylogenetic and bioinformatics analyses. D.A.S and E.J.B did the EM imaging and single particle analyses. T.A.W did most of the *psaL* cloning. N.S. and J.T.N. prepared most of the cell materials. J.T.N carried out the spectral comparison among PCC 7414 PSI oligomers J.W. did the proteomic analyses. M.L. and B.D.B wrote the article, while all other authors have contributed in editing and revising the article.

## Accession numbers

The cloned cyanobacterial *psaL* sequences can be found in GenBank through the following accession numbers: KY575410, KY575411, KY575412, KY575413, KY575414, KY575415, KY575416, KY575417, KY575418, KY575419, KY575420, KY575421, KY575422, KY575423, KY575424. TS-821 Whole Genome Shotgun project has been deposited at DDBJ/ENA/GenBank under the accession MVDI00000000. The version described in this paper is version MVDI01000000.

## Supporting Information

Additional supporting information may be found in the online version of this article.

**Fig. S1.** PsaL antigen design and epitope prediction.

**Fig. S2.** Identification of PSI oligomers in heterocyst-forming cyanobacteria and their unicellular close relatives, contrasted by a few other cyanobacteria.

**Fig. S3.** PAGE analyses of PSI oligomers in cyanobacteria that are evolutionarily distant from heterocyst-forming cyanobacteria, contrasted by a few heterocyst-forming cyanobacteria and close relatives.

**Fig. S4.** Species tree of cyanobacteria and Maximum-likelihood phylogenetic tree of PsaL.

**Fig. S5.** Screenshots of PsaL identification in PSI trimers and tetramers from PCC 7414, PCC 7428 and TS-821.

**Fig. S6.** LOGO plot and alignment of PsaL loop insertion between second and third transmembrane helices (TMH).

**Fig. S7.** Effect of environmental factors on PSI oligomeric states in heterocystforming cyanobacteria and their close relatives.

**Fig. S8.** Macroscopic and microscopic images of strain PCC 7414.

**Fig. S9.** Spectral properties of PSI oligomers in TS-821 under different light intensities.

**Fig. S10.** Absorption spectra of different PSI oligomers from TS-821 and PCC 7414.

**Fig. S11.** HPLC chromatograms of pigment analyses.

**Table S1.** PCR primers and conditions for different psaL cloning.

**Table S2.** Primer sequences used for psaL cloning.

**Dataset S1.** Summary of cyanobacterial PSI oligomeric states, culture conditions, and genomes studied.

