## Supplementary material for "Physiological and Evolutionary Implications of Tetrameric Photosystem I in Cyanobacteria": Fig. S

**This PDF file includes:**

Figs S1 to S11
Tables S1 to S2

**Other supplemental materials for this manuscript include the following:**

Dataset S1

**(a)**

```
>CATSPsaL(Antigen)
MAQAIDASKNRPSDPRNQQVVQPINGDPFIGSLETPISDSGLVKTFIGNLPAYRKGLSPILRGLEVGMAHGYSLYAAT
DPPKPVATVTVPNPPDTFDTTEG
```

**(b)**

```
Antigen   MAQAIDASKNRPSDPRNQQVVQPINGDPFIGSLETPISDSGLVKTFIGNLPAYRKGLSPILRGLEVGMAHGYSLYAATD--PPKP--VATVTVPNPP-DT--FDTTEG
At        ---------AVKSDKTTFQVVQPINGDPFIGSLETPVTSSPLIAWYLSNLPGYRTAVNPLLRGVEVGLAHGFTIYGISSFKEGEPSIAPSLTLTGRKKQPDQLQTADG
T.e.      ---------------MAEELVKPYNGDPFVGHLSTPISDSGLVKTFIGNLPAYRQGLSPILRGLEVGMAHGYAAYGLVSFQKGGS-----------SSDP--LKTSEG
Syn6803   -------------MAESNQVVQAYNGDPFVGHLSTPISDSAFTRTFIGNLPAYRKGLSPILRGLEVGMAHGYSSYGLVTFQGEQG-----------SGDT--LQTADG
TS-821    MAQAIDASKNRPSDPRNQEVVYPSRRDPQIGNLETPINNSSLVKWFINNLPAYRPGITPLRRGLEVGMAHGYSLYAATD--PPKP--VATVTVPNPP-DT--FDTTEG
0.95      -------------------------------------------------------------------------LYAATD--PPKP--VATVTV---------------
0.93      ---------------------------------------------------------------------------------------VATVTVPNPP-DT--FDTT--
0.90      --------------------VQPINGDPFIGSLETP------------------------------------------------------------------------
0.89      -------------------------------------------KTFIGNLPAYRKGLSP-------------------------------------------------
0.87      -----------------------------------------------------------------VGMAHGYSLYAATD--PP-------------------------
0.86      -----DASKNRPSDPRNQQVV---------------------------------------------------------------------------------------
0.75      ----------------------------------------------------YRKGLSPILRGLEVGM----------------------------------------
0.74      ----------------------------------TPISDSGLVKTFIGNL----------------------------------------------------------
0.68      -------------DPRNQQVVQPINGDPF-------------------------------------------------------------------------------
```

**(c)**

```
Antigen   MAQAIDASKNRPSDPRNQQVVQPINGDPFIGSLETPISDSGLVKTFIGNLPAYRKGLSPILRGLEVGMAHGYSLYAATD--PPKP--VATVTVPNPP-DT--FDTTEG
At        ---------AVKSDKTTFQVVQPINGDPFIGSLETPVTSSPLIAWYLSNLPGYRTAVNPLLRGVEVGLAHGFTIYGISSFKEGEPSIAPSLTLTGRKKQPDQLQTADG
T.e.      ---------------MAEELVKPYNGDPFVGHLSTPISDSGLVKTFIGNLPAYRQGLSPILRGLEVGMAHGYAAYGLVSFQKGGS-----------SSDP--LKTSEG
Syn6803   -------------MAESNQVVQAYNGDPFVGHLSTPISDSAFTRTFIGNLPAYRKGLSPILRGLEVGMAHGYSSYGLVTFQGEQG-----------SGDT--LQTADG
TS-821    MAQAIDASKNRPSDPRNQEVVYPSRRDPQIGNLETPINNSSLVKWFINNLPAYRPGITPLRRGLEVGMAHGYSLYAATD--PPKP--VATVTVPNPP-DT--FDTTEG
```

**Fig. S1. PsaL antigen design and epitope prediction**

**(a)** PsaL antigen sequence in fasta format is shown.
**(b)** Alignment of CATSPsaL with PsaL N-termini and loop insertions from *Arabidopsis* (At), *T. elongatus* (T.e.), *Synechocystis* sp. PCC 6803 (Syn6803), and TS-821. First three amino acid residues of loop insertions are in bold format for clarity. The ABCpred epitopes with different scores are aligned as well. Identical fragments to antigen are highlighted in green, and predicted epitopes are underlined.
**(c)** The predicted epitopes using Kolaskar & Tongaonkar method are colored in red.

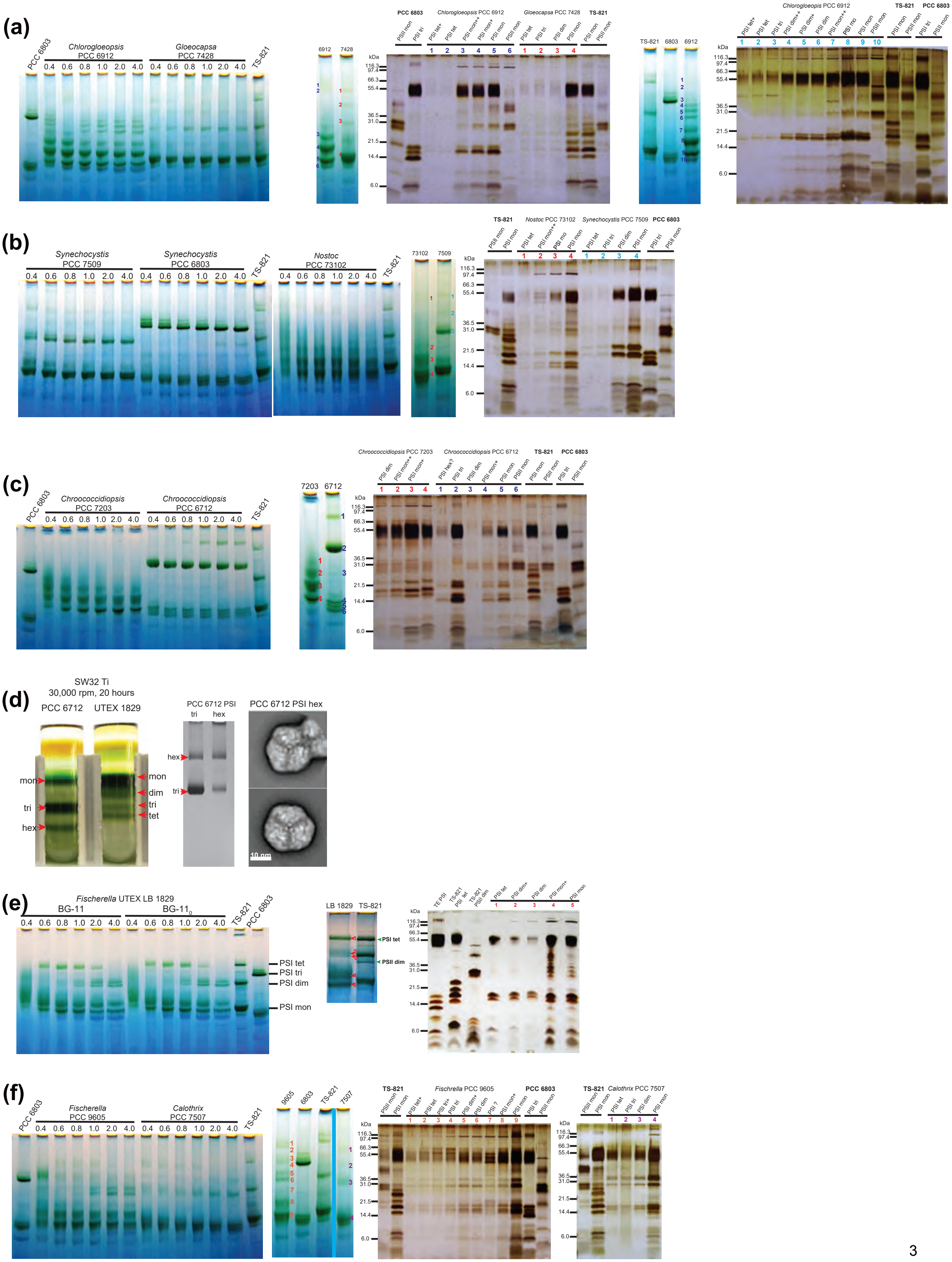

(a)
PCC 6803
Chlorogloeopsis PCC 6912
Gloeocapsa PCC 7428
TS-821
0.4 0.6 0.8 1.0 2.0 4.0
6912
7428
kDa
116.3
97.4
66.3
55.4
36.5
31.0
21.5
14.4
6.0
PSII mon
PSI tri
PSI tet+
PSI tet
PSI mon++
PSI mon+
PSI mon
PSI dim
PSI mon
TS-821
6803
6912
PSI dim++
PSI dim+
PSI mo
(b)
Synechocystis PCC 7509
Synechocystis PCC 6803
Nostoc PCC 73102
73102
7509
(c)
Chroococcidiopsis PCC 7203
Chroococcidiopsis PCC 6712
7203
6712
PSI hex?
(d)
SW32 Ti
30,000 rpm, 20 hours
PCC 6712
UTEX 1829
mon
dim
tri
tet
hex
PCC 6712 PSI
tri
hex
PCC 6712 PSI hex
10 nm
(e)
Fischerella UTEX LB 1829
BG-11
BG-11$_0$
PSI tet
PSI tri
PSI dim
PSI mon
LB 1829
PSII dim
TE PSI
(f)
Fischerella PCC 9605
Calothrix PCC 7507
9605
6803
7507
PSI tri+
PSI ?

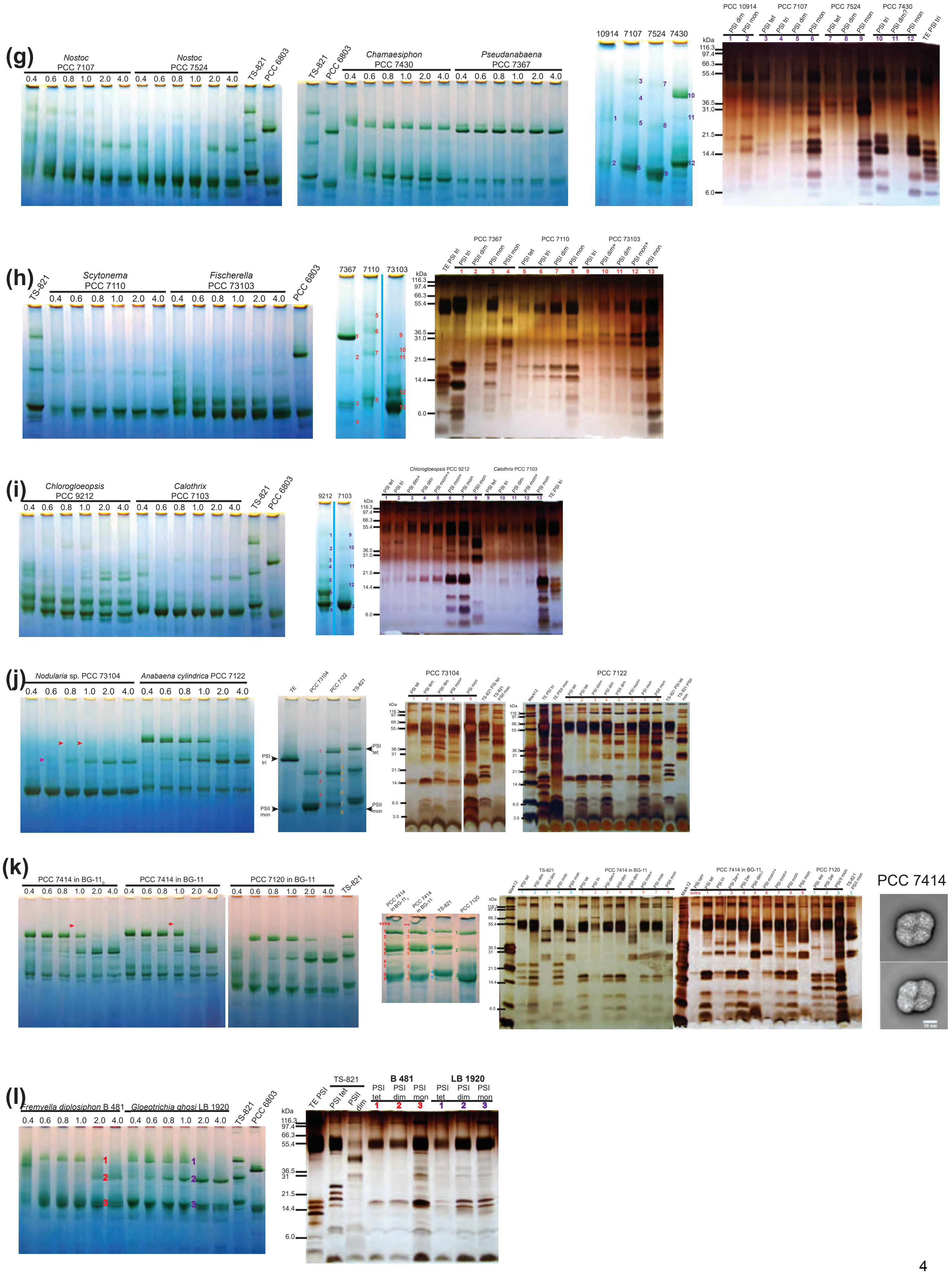
(g)
Nostoc PCC 7107
Nostoc PCC 7524
TS-821
PCC 6803
Chamaesiphon PCC 7430
Pseudanabaena PCC 7367
0.4 0.6 0.8 1.0 2.0 4.0
10914 7107 7524 7430
PCC 10914
PCC 7107
PCC 7524
PCC 7430
kDa
116.3
97.4
66.3
55.4
36.5
31.0
21.5
14.4
6.0
(h)
Scytonema PCC 7110
Fischerella PCC 73103
7367 7110 73103
PCC 7367
PCC 7110
PCC 73103
(i)
Chlorogloeopsis PCC 9212
Calothrix PCC 7103
9212 7103
(j)
Nodularia sp. PCC 73104
Anabaena cylindrica PCC 7122
PSI tri
PSII mon
PSI tet
PCC 73104
PCC 7122
(k)
PCC 7414 in BG-11$_0$
PCC 7414 in BG-11
PCC 7120 in BG-11
PCC 7414
(l)
Fremyella diplosiphon B 481
Gloeotrichia ghosi LB 1920
TE PSI
TS-821
PSI tet
PSII dim
B 481
LB 1920
PSI tet
PSI dim
PSI mon

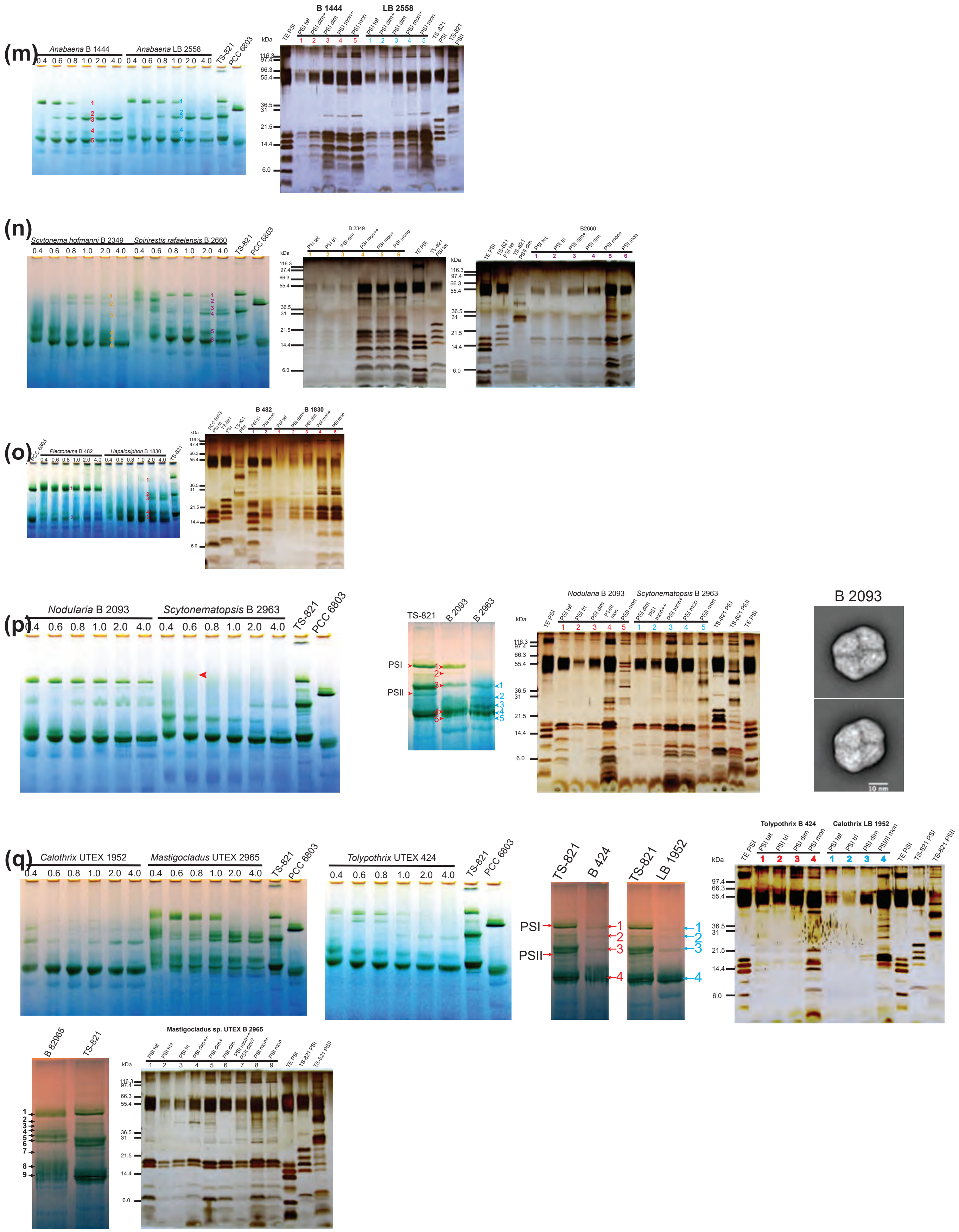
(m)
Anabaena B 1444
Anabaena LB 2558
0.4 0.6 0.8 1.0 2.0 4.0 0.4 0.6 0.8 1.0 2.0 4.0
TS-821
PCC 6803
B 1444
LB 2558
TE PSI
PSI tet
PSI dim+
PSI dim
PSI mon+
PSI mon
kDa
116.3
97.4
66.3
55.4
36.5
31
21.5
14.4
6.0
(n)
Scytonema hofmanni B 2349
Spirirestis rafaelensis B 2660
B 2349
B2660
(o)
Plectonema B 482
Hapalosiphon B 1830
B 482
B 1830
(p)
Nodularia B 2093
Scytonematopsis B 2963
PSI
PSII
B 2093
10 nm
(q)
Calothrix UTEX 1952
Mastigocladus UTEX 2965
Tolypothrix UTEX 424
B 424
LB 1952
Tolypothrix B 424
Calothrix LB 1952
B 82965
Mastigocladus sp. UTEX B 2965

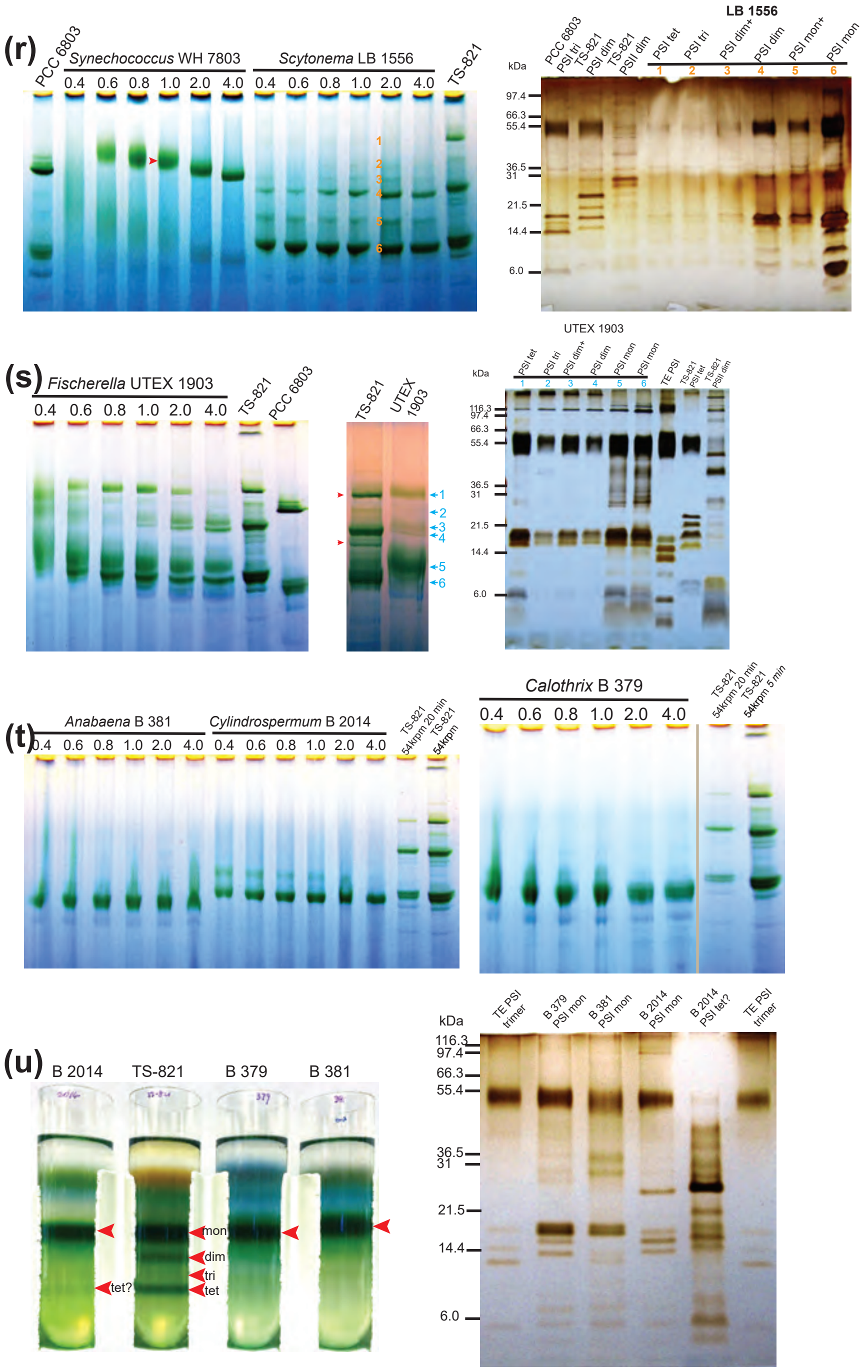


**Fig. S2. Identification of PSI oligomers in heterocyst-forming cyanobacteria and their unicellular close relatives, contrasted by a few other cyanobacteria.**

(Sub-panels from left to right).

**(a)** <u>PAGE analyses of PSI oligomers in PCC 6912 and PCC 7428.</u>

**1st**: BN-PAGE analyses of solubilized thylakoid membrane from PCC 6912 and PCC 7428. Thylakoid membranes with 0.4 mg/mL Chl from PCC 6912 and PCC 7428 were solubilized in different concentrations of DDM (w/v, %), labeled on top of each lane. Solubilized thylakoid membranes from TS-821 and *Synechocystis* sp. PCC 6803 (PCC 6803) with 0.4 mg/mL Chl in 1% DDM were loaded as controls.

~ 2 µg Chl were loaded for each condition. The same solubilization conditions, controls and loading amounts for other strains are used unless specified hereafter. **2nd:** BN-PAGE of solubilized thylakoid membranes PCC 6912 and PCC 7428 for following SDS-PAGE (3rd). The green bands sliced out for second dimension SDS-PAGE (3rd) and PSI identification are numbered. **3rd:** SDS-PAGE of photosystems from PCC 6912 and PCC 7428 for PSI identification. PCC 6803 and TS-821 PSI and PSII from BN-PAGE gel were used as control for PSI identification. Numbered lanes correspond to numbered bands in the 2nd. PSI oligomeric state interpretations were denoted on top of each lane as well. "+" and "++" signs, here and in following figures, denote additional components attached to corresponding oligomers. **4th and 5th:** Additional PAGE analyses of PSI oligomers in PCC 6912 for detailed photosystem profiling. Labeling pattern follows the rules in the 3rd and 4th sub-panels. If not specified, the following panels describing PAGE analyses of PSI oligomers in different strains have similar experimental conditions and labeling system as in panel (a).

**(b)** PAGE analyses of PSI oligomers in PCC 7509 and PCC 73102.

**(c)** PAGE analyses of PCC 7203 and PCC 6712.

**(d)** PCC 6712 trimeric and potential hexameric PSI analyses using SDGC, BN-PAGE, and TEM.

**1st**, SDGC of solubilized thylakoid membrane from PCC 6712. Membranes containing 0.4 mg/mL Chl were solubilized in 2% DDM and 0.6% DDM for PCC 6712 and UTEX LB 1829 respectively. LB 1829 (PAGE analyses in (e)) was used as control. 1 mg total Chl was loaded on each gradient. The position of PSI monomer (mon), dimer (dim), trimer (tri), tetramer (tet) and hexamer (hex) are pointed out by arrows. **2nd**: BN-PAGE of PSI trimer and hexamer isolated from SDGC. The bands corresponding to hexamer and trimer are pointed out. Gel was stained with before imaging. **3rd**: Single particle analysis images of PSI hexamer sample isolated from sucrose density gradient. Two representative species are shown.

**(e)** PAGE analyses of PSI oligomers from LB 1829 cultured in different media.

**(f)** PAGE analyses of PSI oligomers from PCC 9605 and PCC 7507.

**(g)**PAGE analyses of PSI oligomers in PCC 7107, PCC 7524, PCC 7430, PCC 10914 and PCC 7367. The SDS-PAGE of PCC 7367 PSI oligomers is shown in (H). Thylakoid membrane containing 0.2 mg/mL Chl from PCC 10914 was solubilized in 0.6% DDM. Purified *T. elongatus* (TE) PSI trimer was used as control for PSI identification.

**(h)**PAGE analyses of PSI oligomers in PCC 7110, PCC 73103 and PCC 7367.

The shown BN-PAGE of solubilized thylakoid membranes for following SDS-PAGE was loaded with solubilization in 2% DDM and in 0.4% DDM for PCC 7367 and PCC 7110 respectively.

**(i)** PAGE analyses of PSI oligomers in PCC 9212 and PCC 7103.

**(j)** PAGE analyses of PSI oligomers in PCC 73104 and PCC 7122.

**(k)** Structural analyses of PSI oligomers in PCC 7414 and PCC 7120.

PCC 7414 cultured in $BG\text{-}11_0$ and BG-11 were compared. The arrows point at the "extra-large" green band in BN-PAGE. The BN-PAGE of solubilized thylakoid membrane for following SDS-PAGE was loaded with ~ 8 µg Chl for each lane. Last sub-panel shows the single particle analysis images of PSI tetramer isolated from PCC 7414.

**(l)** PAGE analyses for PSI oligomers in B 481 and LB 1920.

**(m)** PAGE analyses of PSI oligomers in B 1444 and LB 2558.

**(n)** PAGE analyses of PSI oligomers in B 2349 and B 2660.

**(o)** PAGE analyses of PSI oligomers in B 482 and B 1830.

**(p)** Structural analyses of PSI oligomers in B 2093 and B 2963 for PSI.

The last sub-panel shows the single particle analysis image of isolated B 2093 PSI tetramer.

**(q)** PAGE analyses of PSI oligomers in LB 1952, B 2965, and B 424.
**(r)** PAGE analysis of PSI oligomeric state in WH 7803 and LB 1556.
The potential PSI trimer from WH 7803 is pointed out by an arrow and analyzed using SDS-PAGE (Fig. S2).
**(s)** PAGE analyses of PSI oligomers in UTEX 1903.
**(t)** BN-PAGE of solubilized thylakoid membranes from B 379, B 381 and B 2014.
Solubilized membranes were centrifuged for 20 min at 54, 000 rpm before loading. TS-821 solubilized membrane with same treatment was used as controls.
**(u)** SDGC and SDS-PAGE analyses of PSI oligomers from B 379, B 381 and B 2014.
Membranes Containing 0.4 mg/mL Chl were solubilized in 0.6% DDM before loading on sucrose gradient. The PSI oligomer positions after centrifugation in the gradient are pointed out for TS-821. Green bands isolated from B 379, arrows point out B 381 and B 2014 for SDS-PAGE analysis too. Potential PSI tetramer in B 2014 is labeled as “tet?”.

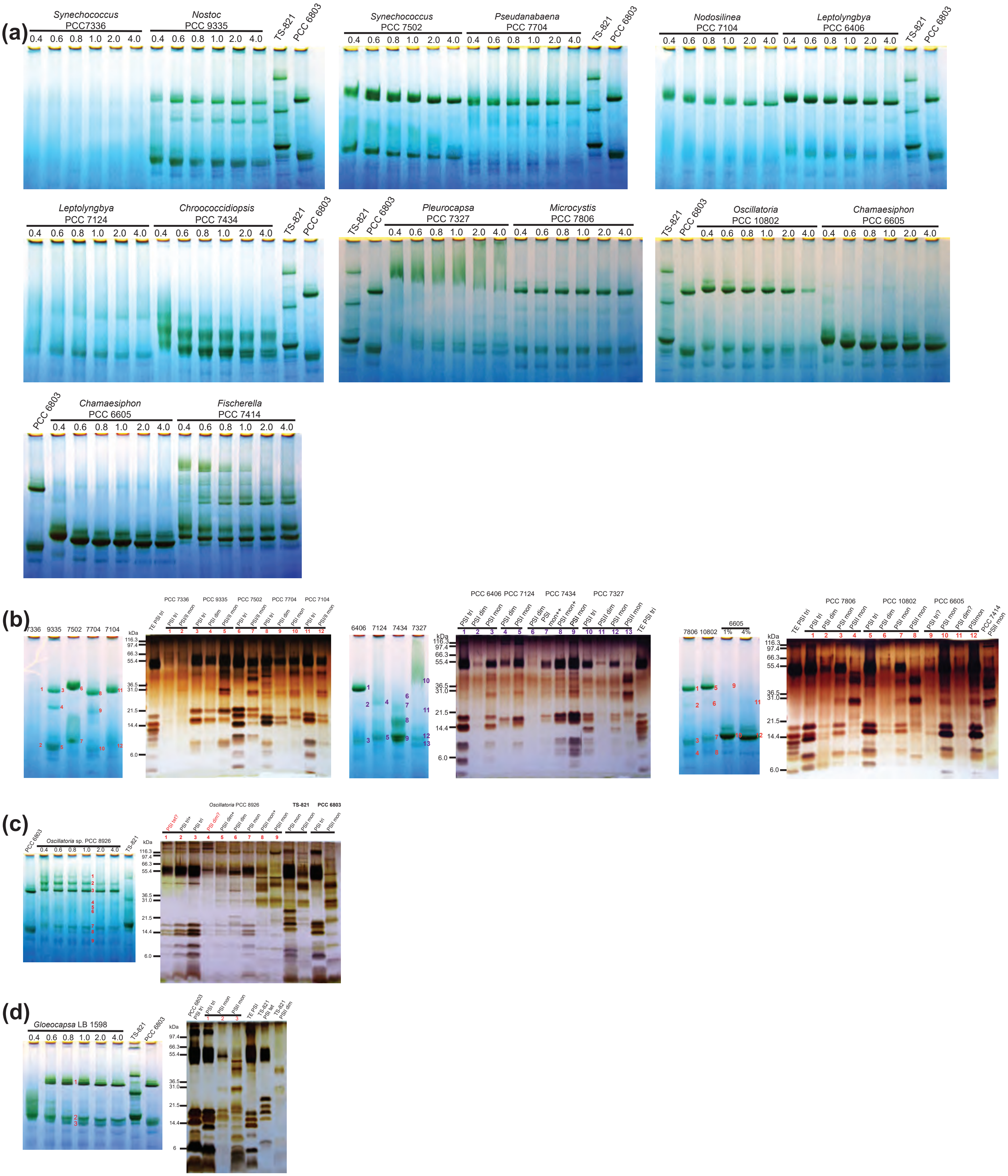
(a)
Synechococcus PCC7336
Nostoc PCC 9335
TS-821
PCC 6803
Synechococcus PCC 7502
Pseudanabaena PCC 7704
Nodosilinea PCC 7104
Leptolyngbya PCC 6406
Leptolyngbya PCC 7124
Chroococcidiopsis PCC 7434
Pleurocapsa PCC 7327
Microcystis PCC 7806
Oscillatoria PCC 10802
Chamaesiphon PCC 6605
Fischerella PCC 7414
0.4 0.6 0.8 1.0 2.0 4.0
(b)
kDa
116.3 97.4 66.3 55.4 36.5 31.0 21.5 14.4 6.0
(c)
Oscillatoria sp. PCC 8926
Oscillatoria PCC 8926
(d)
Gloeocapsa LB 1598

(e)

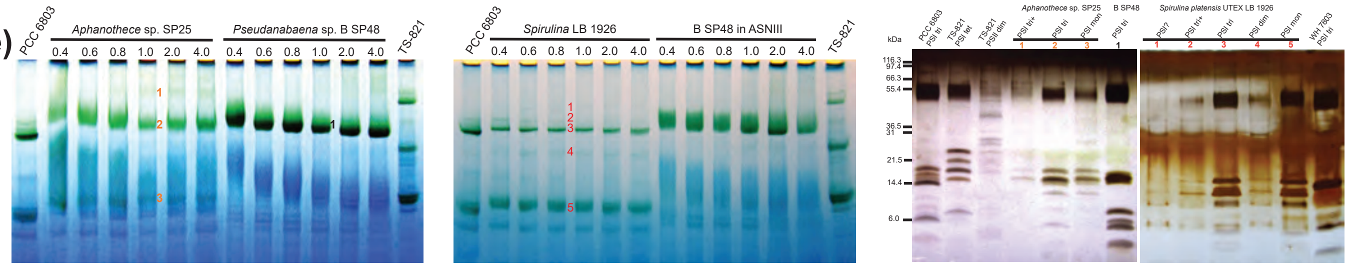


**Fig. S3. PAGE analyses of PSI oligomers in cyanobacteria that are evolutionarily distant from heterocyst-forming cyanobacteria, contrasted by a few heterocyst-forming cyanobacteria and close relatives.**

**(a)** BN-PAGE of solubilized thylakoid membrane from PCC 7336, PCC 9335, PCC 7502, PCC 7704, PCC 7104, PCC 6406, PCC 7124, PCC 7434, 7327, PCC 7806, PCC 10802, and PCC 6605. ~ 2 μg Chl were loaded for each solubilization condition except that ~3.2 μg Chl were loaded for each lane in last sub-panel where PCC 6605 is contrasted by PCC 7414.

**(b)** PAGE analyses for PSI oligomer identification. Strains are as listed in (a). For PCC 6605, two solubilization conditions (1% DDM and 4% DDM) were used, with ~3.2 μg Chl loaded for each lane.

**(c)** PAGE analysis of PSI complexes in PCC 8926. “?” denotes the uncertainty of the oligomeric states.

**(d)** PAGE analysis of PSI oligomeric state in LB 1598.

**(e)** PAGE analyses of PSI oligomers of SP25, B SP48, LB 1926 and WH 7803. B SP48 grown in modified ASNIII medium and BG-11 were compared in BN-PAGE. WH 7803 PSI band was sliced from the BN-PAGE gel, as shown in Fig. S2r.

The experimental conditions for PAGE analyses and labeling system is the same as Fig. S2 if not specified.

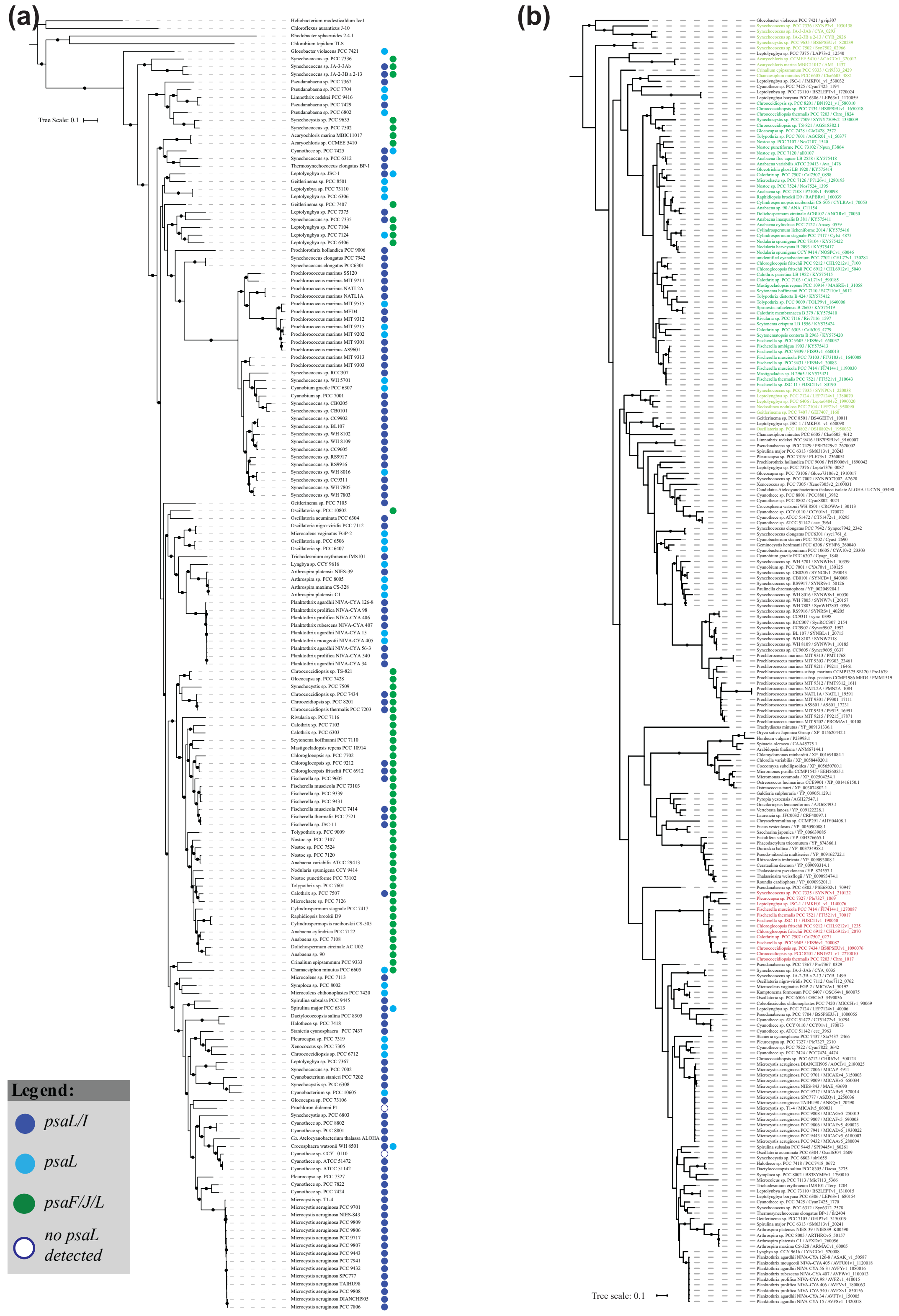


**Fig. S4. Species tree of cyanobacteria and Maximum-likelihood phylogenetic tree of PsaL.**

**(a)** Species tree of cyanobacteria. The *psaL* gene is color coded according to its location in relation to *psaF/J* and *psaI*, and denoted after each strain. Bootstrap values ≥ 70% are indicated by black dots.
**(b)** Maximum-likelihood phylogenetic tree of PsaL. PsaL encoded by *psaL* located downstream of *psaF/psaJ* are colored in light and dark green, with dark green denoting the heterocyst-forming species and their close relatives. Potential far-red light responsive PsaL are colored in red.

### *Fischerella* PCC 7414

**PSI trimer**

| | | Score | Mass | Matches | Sequences | emPAI | |
|---|---|---|---|---|---|---|---|
| 6.1 | tr\|G6FY38\|G6FY38_9C | 460 | 18317 | 10 (10) | 3 (3) | 1.15 | Photosystem I reaction center subunit XI OS=Fischerella sp. JSC-11 GN |

| Query | Dupes | Observed | Mr(expt) | Mr(calc) | ppm | M | Score | Expect | Rank | U | Peptide |
|---|---|---|---|---|---|---|---|---|---|---|---|
| 2827 | ▶6 | 949.5008 | 1896.9870 | 1896.9792 | 4.10 | 0 | 114 | 7.4e-10 | ▶1 | U | R.DPQIGNLETPINSSALTK.W |
| 2995 | | 660.6687 | 1978.9843 | 1978.9764 | 3.99 | 0 | 21 | 0.03 | ▶1 | | R.GLEVGMAHGYWIFGPFAK.L |
| 3021 | | 666.0004 | 1994.9795 | 1994.9713 | 4.09 | 0 | 23 | 0.044 | ▶1 | | R.GLEVGMAHGYWIFGPFAK.L + Oxidation (M) |
| 3125 | | 688.0420 | 2061.1041 | 2061.0948 | 4.52 | 0 | 35 | 0.016 | ▶1 | U | K.WFINNLPAYRPGITPFR.R |

**PSI tetramer**

| | | Score | Mass | Matches | Sequences | emPAI | |
|---|---|---|---|---|---|---|---|
| 7.1 | tr\|G6FY38\|G6FY38_9C | 536 | 18317 | 11 (11) | 3 (3) | 1.60 | Photosystem I reaction center subunit XI OS=Fischerella sp. JSC-11 GN |

| Query | Dupes | Observed | Mr(expt) | Mr(calc) | ppm | M | Score | Expect | Rank | U | Peptide |
|---|---|---|---|---|---|---|---|---|---|---|---|
| 3589 | ▶6 | 949.5001 | 1896.9856 | 1896.9792 | 3.33 | 0 | 147 | 3.3e-13 | ▶1 | U | R.DPQIGNLETPINSSALTK.W |
| 3797 | | 660.6687 | 1978.9843 | 1978.9764 | 3.99 | 0 | 26 | 0.024 | ▶1 | | R.GLEVGMAHGYWIFGPFAK.L |
| 3798 | | 990.5018 | 1978.9890 | 1978.9764 | 6.37 | 0 | 46 | 8.5e-05 | ▶1 | | R.GLEVGMAHGYWIFGPFAK.L |
| 3838 | | 666.0001 | 1994.9784 | 1994.9713 | 3.54 | 0 | 28 | 0.0053 | ▶1 | | R.GLEVGMAHGYWIFGPFAK.L + Oxidation (M) |
| 3977 | | 688.0399 | 2061.0979 | 2061.0948 | 1.50 | 0 | 35 | 0.011 | ▶1 | U | K.WFINNLPAYRPGITPFR.R |

### *Gloeocapsa* PCC 7428

**PSI trimer**

| | | Score | Mass | Matches | Sequences | emPAI | |
|---|---|---|---|---|---|---|---|
| 9.1 | tr\|K9XF28\|K9XF28_9C | 676 | 18498 | 65 (40) | 9 (6) | 4.48 | Photosystem I reaction center subunit XI OS=Gloeocapsa sp. PCC 7428 |

| Query | Dupes | Observed | Mr(expt) | Mr(calc) | ppm | M | Score | Expect | Rank | U | Peptide |
|---|---|---|---|---|---|---|---|---|---|---|---|
| 105 | ▶4 | 421.2186 | 840.4226 | 840.4202 | 2.84 | 0 | 12 | 2 | ▶4 | | K.NRPSDPR.N |
| 818 | ▶7 | 532.2763 | 1062.5381 | 1062.5346 | 3.29 | 0 | 56 | 0.00044 | ▶1 | U | R.NQEVVYPSK.R |
| 1513 | ▶13 | 407.2205 | 1218.6397 | 1218.6357 | 3.34 | 1 | 41 | 0.016 | ▶1 | U | R.NQEVVYPSKR.D |
| 1532 | ▶15 | 610.3284 | 1218.6423 | 1218.6357 | 5.45 | 1 | 44 | 0.0012 | ▶1 | U | R.NQEVVYPSKR.D |
| 3345 | | 629.3253 | 1884.9539 | 1884.9442 | 5.17 | 1 | 7 | 0.72 | ▶1 | U | K.NRPSDPRNQEVVYPSK.R |
| 3409 | | 959.4910 | 1916.9674 | 1916.9607 | 3.48 | 0 | 77 | 2.7e-07 | ▶1 | U | R.GLEAGMAHGYWLLGPFAK.L |
| 3410 | ▶1 | 639.9965 | 1916.9676 | 1916.9607 | 3.57 | 0 | 29 | 0.037 | ▶1 | U | R.GLEAGMAHGYWLLGPFAK.L |
| 3429 | ▶1 | 967.4882 | 1932.9618 | 1932.9556 | 3.18 | 0 | 32 | 0.0077 | ▶1 | U | R.GLEAGMAHGYWLLGPFAK.L + Oxidation (M) |
| 3430 | ▶1 | 645.3282 | 1932.9627 | 1932.9556 | 3.67 | 0 | 19 | 0.086 | ▶1 | U | R.GLEAGMAHGYWLLGPFAK.L + Oxidation (M) |
| 3441 | ▶4 | 970.0132 | 1938.0118 | 1938.0058 | 3.11 | 0 | 102 | 8.3e-09 | ▶1 | | R.DPQIGNLETPINNSSLVK.W |
| 3602 | ▶1 | 676.7126 | 2027.1159 | 2027.1105 | 2.69 | 0 | 45 | 0.0039 | ▶1 | | K.WFINNLPAYRPGITPLR.R |
| 3603 | | 1014.5681 | 2027.1216 | 2027.1105 | 5.47 | 0 | 11 | 0.19 | ▶1 | | K.WFINNLPAYRPGITPLR.R |
| 3623 | | 511.2704 | 2041.0524 | 2041.0453 | 3.50 | 2 | 5 | 1.5 | ▶5 | U | K.NRPSDPRNQEVVYPSKR.D |
| 3756 | ▶2 | 699.0450 | 2094.1131 | 2094.1069 | 2.97 | 1 | 21 | 0.52 | ▶1 | | K.RDPQIGNLETPINNSSLVK.W |
| 3758 | ▶1 | 1048.0647 | 2094.1148 | 2094.1069 | 3.79 | 1 | 50 | 4.3e-05 | ▶1 | | K.RDPQIGNLETPINNSSLVK.W |

**PSI tetramer**

| | | Score | Mass | Matches | Sequences | emPAI | |
|---|---|---|---|---|---|---|---|
| 7.1 | tr\|K9XF28\|K9XF28_9C | 905 | 18498 | 58 (46) | 7 (6) | 3.54 | Photosystem I reaction center subunit XI OS=Gloeocapsa sp. PCC 742 |

| Query | Dupes | Observed | Mr(expt) | Mr(calc) | ppm | M | Score | Expect | Rank | U | Peptide |
|---|---|---|---|---|---|---|---|---|---|---|---|
| 841 | ▶25 | 532.2765 | 1062.5384 | 1062.5346 | 3.64 | 0 | 56 | 0.00038 | ▶1 | U | R.NQEVVYPSK.R |
| 1579 | ▶10 | 407.2206 | 1218.6400 | 1218.6357 | 3.56 | 1 | 35 | 0.015 | ▶1 | U | R.NQEVVYPSKR.D |
| 1582 | ▶7 | 610.3276 | 1218.6406 | 1218.6357 | 4.05 | 1 | 46 | 0.0024 | ▶1 | U | R.NQEVVYPSKR.D |
| 3501 | | 629.3238 | 1884.9495 | 1884.9442 | 2.84 | 1 | 7 | 1.2 | ▶2 | U | K.NRPSDPRNQEVVYPSK.R |
| 3590 | | 967.4847 | 1932.9549 | 1932.9556 | -0.36 | 0 | 22 | 0.049 | ▶1 | U | R.GLEAGMAHGYWLLGPFAK.L + Oxidation (M) |
| 3591 | | 645.3297 | 1932.9671 | 1932.9556 | 5.95 | 0 | 18 | 0.24 | ▶1 | U | R.GLEAGMAHGYWLLGPFAK.L + Oxidation (M) |
| 3598 | ▶3 | 970.0129 | 1938.0112 | 1938.0058 | 2.79 | 0 | 95 | 5e-09 | ▶1 | | R.DPQIGNLETPINNSSLVK.W |
| 3753 | | 676.7129 | 2027.1168 | 2027.1105 | 3.14 | 0 | 42 | 0.0043 | ▶1 | | K.WFINNLPAYRPGITPLR.R |
| 3916 | ▶2 | 699.0453 | 2094.1142 | 2094.1069 | 3.49 | 1 | 35 | 0.0026 | ▶1 | | K.RDPQIGNLETPINNSSLVK.W |
| 3918 | ▶1 | 1048.0666 | 2094.1187 | 2094.1069 | 5.66 | 1 | 84 | 3.2e-08 | ▶1 | | K.RDPQIGNLETPINNSSLVK.W |

### *Chroococcidiopsis* TS-821

**PSI trimer**

| | | Score | Mass | Matches | Sequences | emPAI | |
|---|---|---|---|---|---|---|---|
| 9.1 | tr\|V5JYE0\|V5JYE0_9C | 486 | 18478 | 38 (18) | 7 (5) | 2.13 | Photosystem I reaction center subunit XI OS=Chroococcidiopsis sp. |

| Query | Dupes | Observed | Mr(expt) | Mr(calc) | ppm | M | Score | Expect | Rank | U | Peptide |
|---|---|---|---|---|---|---|---|---|---|---|---|
| 103 | ▶5 | 421.2184 | 840.4223 | 840.4202 | 2.47 | 0 | 15 | 0.7 | ▶1 | | K.NRPSDPR.N |
| 978 | ▶5 | 546.2767 | 1090.5389 | 1090.5407 | -1.65 | 0 | 60 | 4.9e-05 | ▶1 | U | R.NQEVVYPSR.R |
| 1820 | | 624.3261 | 1246.6375 | 1246.6418 | -3.42 | 1 | 16 | 1.2 | ▶2 | U | R.NQEVVYPSRR.D |
| 1825 | ▶13 | 416.5542 | 1246.6408 | 1246.6418 | -0.84 | 1 | 35 | 0.03 | ▶1 | U | R.NQEVVYPSRR.D |
| 3916 | ▶2 | 970.0112 | 1938.0078 | 1938.0058 | 1.03 | 0 | 104 | 1e-09 | ▶1 | U | R.DPQIGNLETPINNSSLVK.W |
| 3941 | | 649.3349 | 1944.9829 | 1944.9920 | -4.70 | 0 | 16 | 1.4 | ▶2 | U | R.GLEVGMAHGYWLLGPFAK.L |
| 3962 | | 654.6694 | 1960.9863 | 1960.9869 | -0.33 | 0 | 25 | 0.055 | ▶1 | U | R.GLEVGMAHGYWLLGPFAK.L + Oxidation (M) |
| 4063 | | 676.7109 | 2027.1108 | 2027.1105 | 0.16 | 0 | 25 | 0.018 | ▶1 | U | K.WFINNLPAYRPGITPLR.R |
| 4262 | ▶2 | 699.0425 | 2094.1056 | 2094.1069 | -0.62 | 1 | 44 | 0.0018 | ▶1 | U | R.RDPQIGNLETPINNSSLVK.W |
| 4263 | ▶1 | 1048.0602 | 2094.1058 | 2094.1069 | -0.52 | 1 | 84 | 2.1e-08 | ▶1 | U | R.RDPQIGNLETPINNSSLVK.W |

**PSI tetramer**

| | | Score | Mass | Matches | Sequences | emPAI | |
|---|---|---|---|---|---|---|---|
| 6.1 | tr\|V5JYE0\|V5JYE0_9C | 647 | 18478 | 42 (22) | 8 (6) | 3.58 | Photosystem I reaction center subunit XI OS=Chroococcidiopsis sp. TS |

| Query | Dupes | Observed | Mr(expt) | Mr(calc) | ppm | M | Score | Expect | Rank | U | Peptide |
|---|---|---|---|---|---|---|---|---|---|---|---|
| 104 | ▶9 | 421.2181 | 840.4217 | 840.4202 | 1.82 | 0 | 18 | 0.78 | ▶3 | | K.NRPSDPR.N |
| 1082 | ▶6 | 546.2774 | 1090.5403 | 1090.5407 | -0.41 | 0 | 54 | 5.9e-05 | ▶1 | U | R.NQEVVYPSR.R |
| 1945 | ▶5 | 416.5547 | 1246.6423 | 1246.6418 | 0.41 | 1 | 34 | 0.017 | ▶1 | U | R.NQEVVYPSRR.D |
| 3805 | | 638.6571 | 1912.9495 | 1912.9503 | -0.45 | 1 | 11 | 0.64 | ▶1 | U | K.NRPSDPRNQEVVYPSR.R |
| 3855 | ▶4 | 970.0111 | 1938.0077 | 1938.0058 | 0.96 | 0 | 89 | 2.5e-08 | ▶1 | | R.DPQIGNLETPINNSSLVK.W |
| 3882 | | 649.3397 | 1944.9972 | 1944.9920 | 2.64 | 0 | 36 | 0.017 | ▶1 | U | R.GLEVGMAHGYWLLGPFAK.L |
| 3903 | ▶1 | 654.6705 | 1960.9896 | 1960.9869 | 1.35 | 0 | 34 | 0.0019 | ▶1 | U | R.GLEVGMAHGYWLLGPFAK.L + Oxidation (M) |
| 4023 | ▶1 | 676.7110 | 2027.1112 | 2027.1105 | 0.34 | 0 | 45 | 0.00021 | ▶1 | | K.WFINNLPAYRPGITPLR.R |
| 4195 | ▶2 | 1048.0613 | 2094.1080 | 2094.1069 | 0.53 | 1 | 84 | 1.2e-08 | ▶1 | | R.RDPQIGNLETPINNSSLVK.W |
| 4197 | ▶4 | 699.0435 | 2094.1087 | 2094.1069 | 0.87 | 1 | 59 | 4.5e-05 | ▶1 | | R.RDPQIGNLETPINNSSLVK.W |

**Fig. S5. Screenshots of PsaL identification in PSI trimers and tetramers from PCC 7414, PCC 7428 and TS-821.**

**(a)**

LOGO plot of linker sequence between TMH2 and TMH3 of PsaL in tet/dim/mon PSI

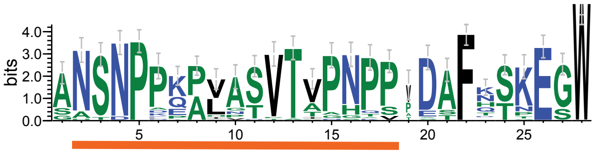


| | |
|---|---|
| TS-821 | AATDPPKPVATVTVPNPP-DTFDTTEGW |
| PCC 7428 | AATDPPKPVATVTVPNPP-DTFDTTEGW |
| PCC 7509 | ANSNPPQPVANVTTANKVPDAFKSPEGW |
| PCC 7203 | SASNPPQPVATTTTGGQVPSTFKSPESW |
| PCC 6912 | ANSNPPKPVNSVTIPNPP-EAFQSNEGW |
| PCC 9212 | ANSNPPKPVNSVTIPNPP-EAFQSNEGW |
| PCC 73102 | SNSNPPKALPSVTVPNPPVDAFNSKESW |
| PCC 9605 | ANSNPPQPVASVTVPNTP-DAFQSKEGW |
| PCC 10914 | GNSNPPQPHVTVTTPNPP-DAFRSKEGW |
| PCC 73103 | ANSNPPEPVASVTAPHPS-DAFHTKEGW |
| PCC 7107 | ANSNPAKALNSVTGASAP-DAFLSKEGW |
| PCC 7524 | ANSNPPKALASVTVPNTP-DAFSSKEGW |
| PCC 7110 | ANSNPPQPHVTVTTPNPP-DAFKSGEGW |
| PCC 7103 | ANSNPKPALGSVTVPNPP-DAFTSKEGW |
| PCC 7414 | ANSNPPEPVASVTAPHPS-DAFHTKEGW |
| PCC 7122 | SNSNPDKALASVTVPNPPVDAFNSKESW |
| PCC 7120 | ANSNPPTALASVTVPNPP-DAFQSKEGW |
| UTEX B 481 | SHSNPPKALASVTAGNPP-DAFTSNEGW |
| ATCC 29413 | ANSNPPKALASVTVPNPP-DAFQSKEGW |
| PCC 9009 | GNSNPLPPVPTVTVPNPP-DSFKTKEGW |
| PCC 9339 | ANSNPPEPVASVTAPHPS-EAFHTKEGW |
| PCC 9431 | ANSNPPEPVASVTAPHPS-DAFHTKEGW |
| PCC 7507 | ANSNPPKALASVTVPNPPADAFNSKEGW |
| PCC 7434 | SASNPPQPVATVTTNGQVPATFKSPESW |
| UTEX 1903 | ANSNPPEPVASVTAPHPS-EAFHTKEGW |
| UTEX B 381 | SNSNPDKALASVTVPNPPVDAFNSKESW |
| UTEX B 2014 | ANSNPPKALPSVTVPNPPTDAFNSKESW |
| UTEX B 2965 | ANSNPPEPVASVTAPHPS-DAFHTKEGW |
| PCC 73104 | ANSNPAKALASVTVPNPPVDAFNSKESW |
| UTEX B 2093 | ANSNPEKALASVTVPNPPVDAFNSKESW |
| UTEX LB 1952 | ANSNPKPALGSVTVPNPP-DAFTSKEGW |
| UTEX B 379 | GNSNPPQPAATVTVPNPP-DSFKTKEGW |
| UTEX LB 2558 | ANSNPPKALASVTVPNPP-DAFQSKEGW |
| UTEX B 2963 | ANSNPNKAIGSPAAPNPP-DAFTSKEGW |
| UTEX B 424 | ANSNPPQPVATVTVPNPP-DSFKTKEGW |
| UTEX B 2660 | GNSNPPQPVATVTVPNPP-DSFKTKEGW |
| UTEX LB 1920 | ANSNPPKAIASVTVPNPP-DAFNSKEGW |
| UTEX LB 1556 | ANSNPPQPVASVTVPNPP-DAFKSNEGW |

**(b)**

LOGO plot of linker sequence between TMH2 and TMH3 of PsaL in tet/dim/mon PSI

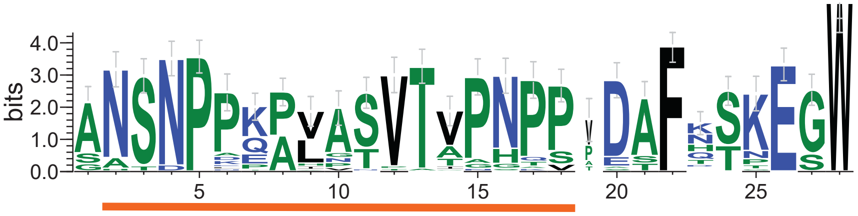


```
PCC 6605(FJL) ASVNPAAPLPSVTVPTPP-NNIKDGKGW
PCC 6605(L)   AATNPPPPIATLTVPNPP-DVFASTQGW
PCC 6308      GAVGVSKPTETLTTPNVPMDLA-TKEGW
WH 7803       AGNGPNVQPADATIDNPPADLF-TKAGW
PCC 7104      SGAGVNTEVTKTTTPFTPPASLETDEGW
PCC 6406      SGAGISKAVTEATAPYTPPAAFSTNEGW
PCC 10802     GSVGVQKPVATITTPNPP-EGLGTDEGW
PCC 7124(FJL) SGAGVNSAVAKSTAPFSPPEALSHDEGW
PCC 7124(L)   GLVSLKPAADL---P-PDTDPLKTSEGW
PCC 7336      SYVT-DD----------GDVFIGSKEGW
PCC 7502      AYSQPRK-----------DDWLGGENGW
PCC 7704      GRFSRKSAAESP----AGTAPEFTSEGW
PCC 7327(*)   GLVSFQGGRAIG----TGDASLKSSEGW
PCC 7806      GLVSFQGKAASG-------DPLQSSEGW
PCC 7367      GIVSYQGDKAGT----PG--TLKSADGW
PCC 6712      GIATFQDKPTNS------EDSLQSSEGW
```

**Fig. S6. LOGO plot and alignment of PsaL loop insertion between second and third transmembrane helices (TMH).**

The LOGO plot of linker sequence between PsaL TMH2 and TMH3 from studied heterocyst-forming cyanobacteria and their close relatives showed unique highly conserved sequence. The unique part of the sequence is underlined. Corresponding PsaL motifs from studied HCR in *PsaF/J/L* **(a)** and other studied strains **(b)** are aligned for comparison. The multi-proline regions of the PsaLs from HCR and PCC 6605 are colored according to the LOGO plot. FJL/L in the brackets denotes the *psaL* gene localization in relation to *psaF/J*. "(*)", the PsaL sequence (Ple_2310, Fig. S4b) from PCC 7327 that is not potentially far-red light responsive was used.

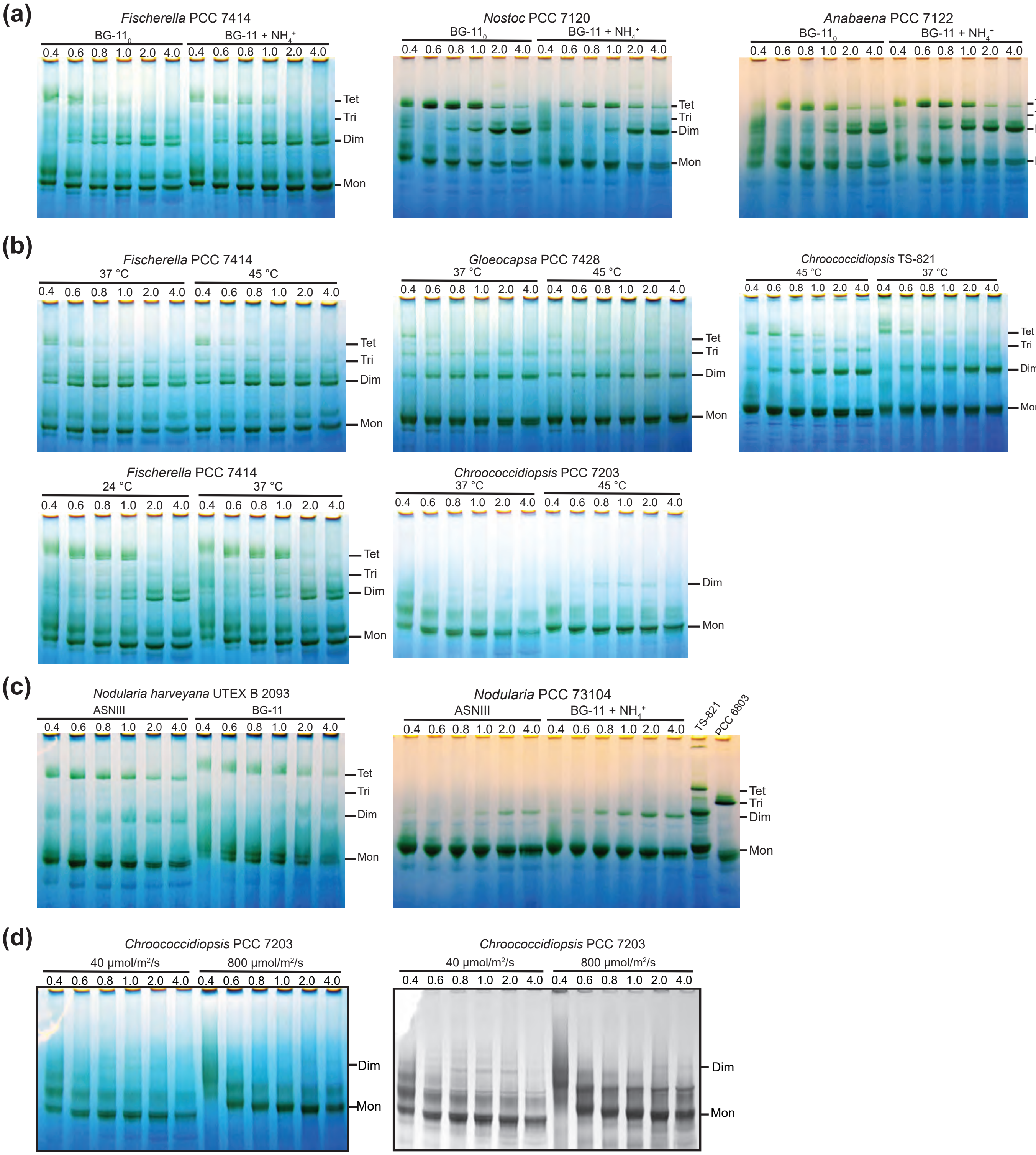


**Fig. S7. Effect of environmental factors on PSI oligomeric states in heterocyst-forming cyanobacteria and their close relatives.**

**(a)** Effect of nitrogen source on PSI oligomeric states in heterocyst-forming cyanobacteria.

**(b)** Effect of temperature on PSI oligomeric states in heterocyst-forming cyanobacteria and their close relatives

**(c)** Effect of salinity on PSI tetramer formation in heterocyst-forming cyanobacteria. Note that the experiment on PCC 73104 also has difference in nitrogen source.

**(d)** Comparison of PCC 7203 PSI oligomeric profiles under different light intensities.

For PCC 7414, PCC 7428 and PCC 7203, when comparing 37 °C with 45 °C (panel (B)), thylakoid membranes with 0.2 mg/mL Chl from each condition were solubilized in different concentrations of DDM (w/v, %), labeled on top of each lane. ~ 1.6 μg Chl were loaded for each condition. This is also true for panel (d). Other experiments follow the general conditions as described in Fig. S2.

(a) (b) (c)

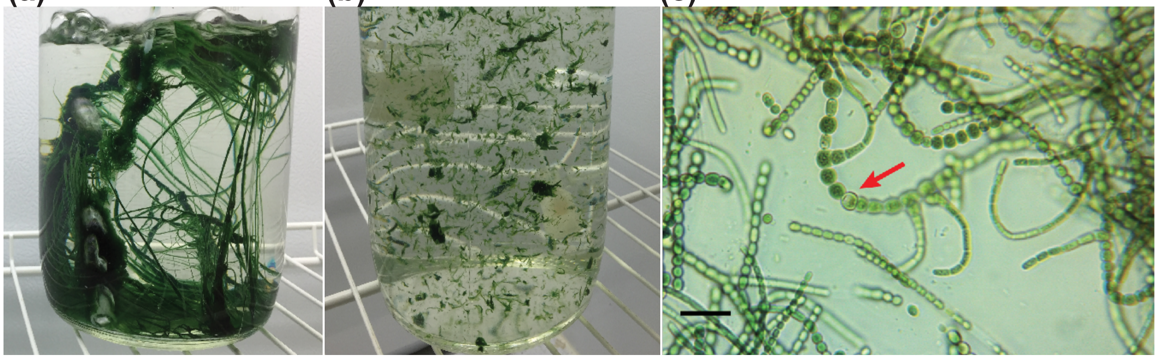

**Fig. S8. Macroscopic and microscopic images of strain PCC 7414**

**(a)** Macroscopic image shows the PCC 7414 cell aggregation into large bundles.

**(b)** Those bundles are broken into smaller but significant clumps for inoculation.

**(c)** Microscopic image shows a typical intertwined cell aggregation in PCC 7414 culture. A heterocyst is pointed out by an arrow. Scale bar, 20 µm.

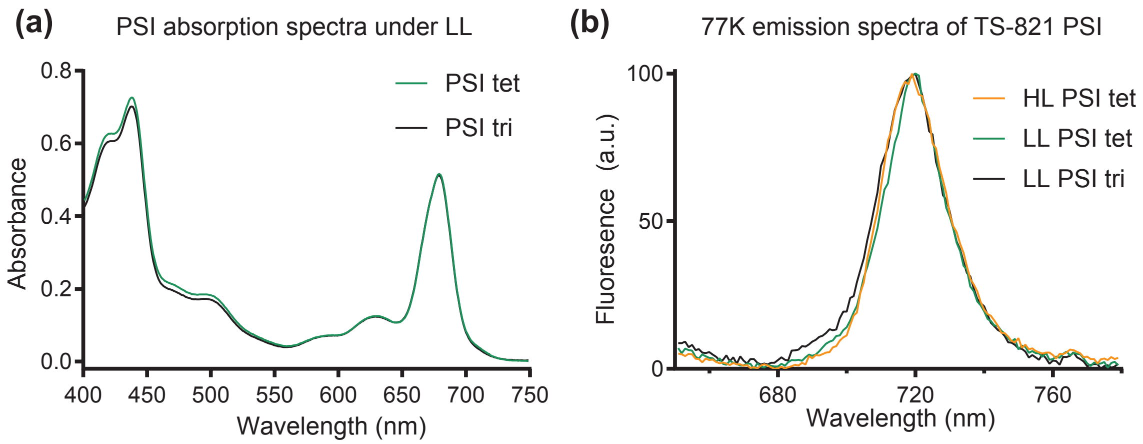


**Fig. S9. Spectral properties of PSI oligomers in TS-821 under different light intensities.** High light (HL) and low light (LL) conditions are described as in materials and methods.

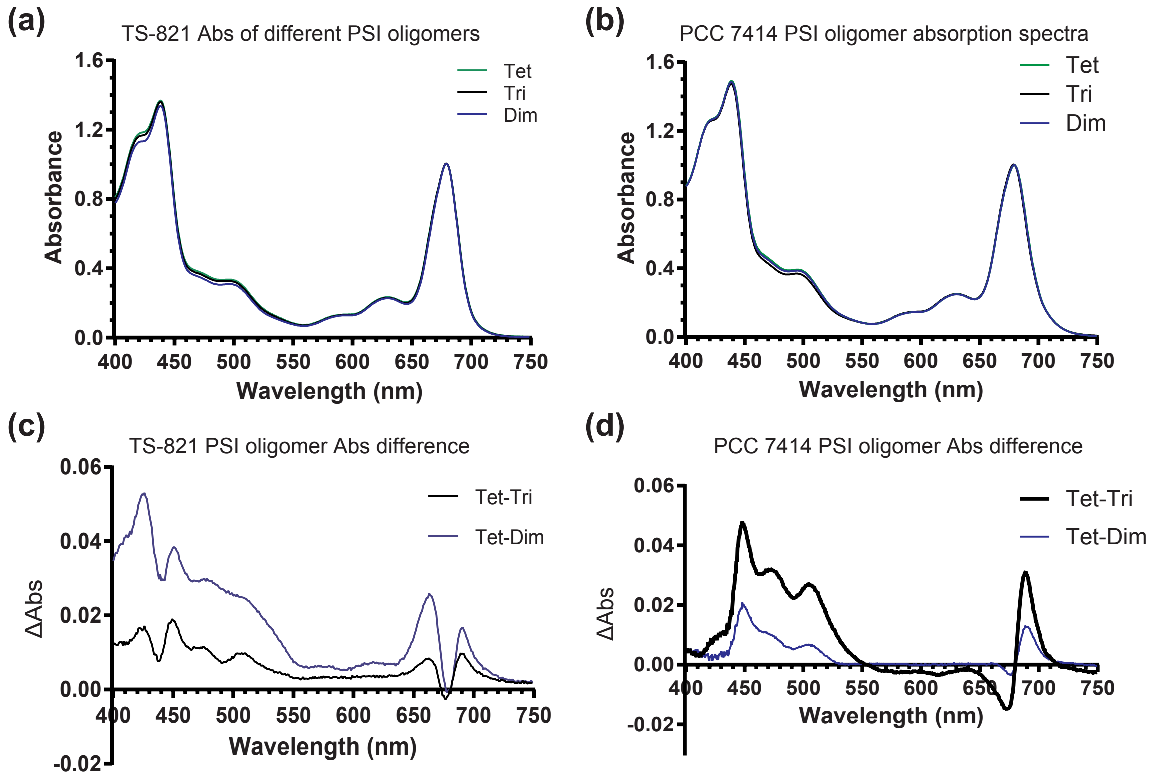


**Fig. S10. Absorption spectra of different PSI oligomers from TS-821 and PCC 7414.**

**(a)** Absorption spectra of different PSI oligomers isolated from TS-821, using SDGC. The centrifugation result for PSI oligomer isolation is shown in Fig. 4E.

**(b)** Absorption spectra of different PSI oligomers isolated from PCC 7414, using SDGC.

**(c)** Absorption difference spectra between different oligomers, calculated from (A).

**(d)** Absorption difference spectra between different oligomers, calculated from (B).

**(a)**

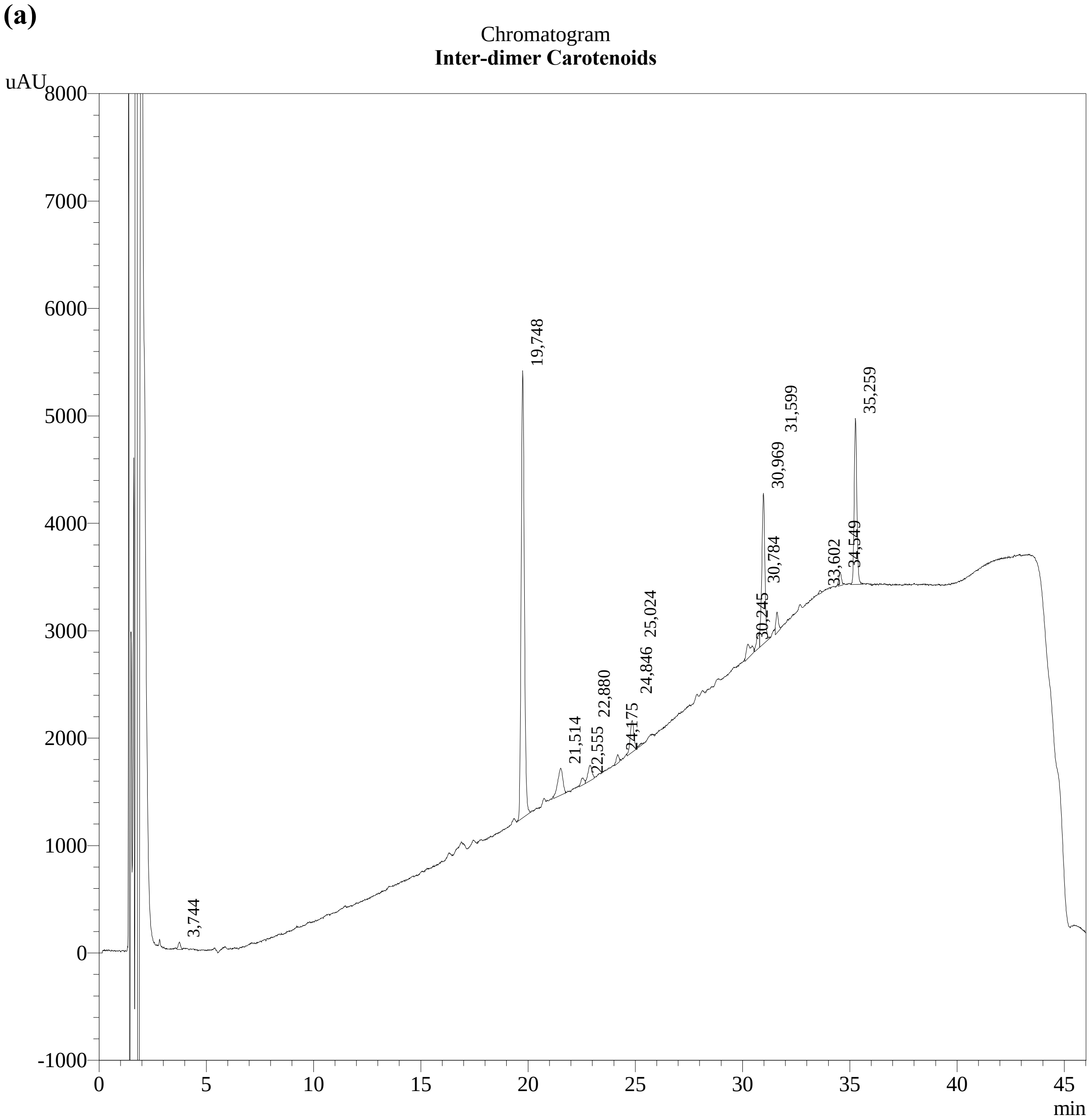


PeakTable

PDA Ch1 450nm

| Peak# | Name | Ret. Time | Area |
|---|---|---|---|
| 1 | unknown | 3,744 | 506 |
| 2 | myxo like | 19,748 | 42777 |
| 3 | unknown | 21,514 | 4175 |
| 5 | unknown | 22,880 | 1935 |
| 7 | canta | 24,846 | 3540 |
| 9 | unknown | 30,245 | 2020 |
| 10 | chla | 30,784 | 1299 |
| 11 | echi | 30,969 | 13784 |
| 12 | chla | 31,599 | 1523 |
| 13 | pht | 33,602 | 184 |
| 14 | unknown | 34,549 | 1038 |
| 15 | beta carotene | 35,259 | 13169 |

**(b)**

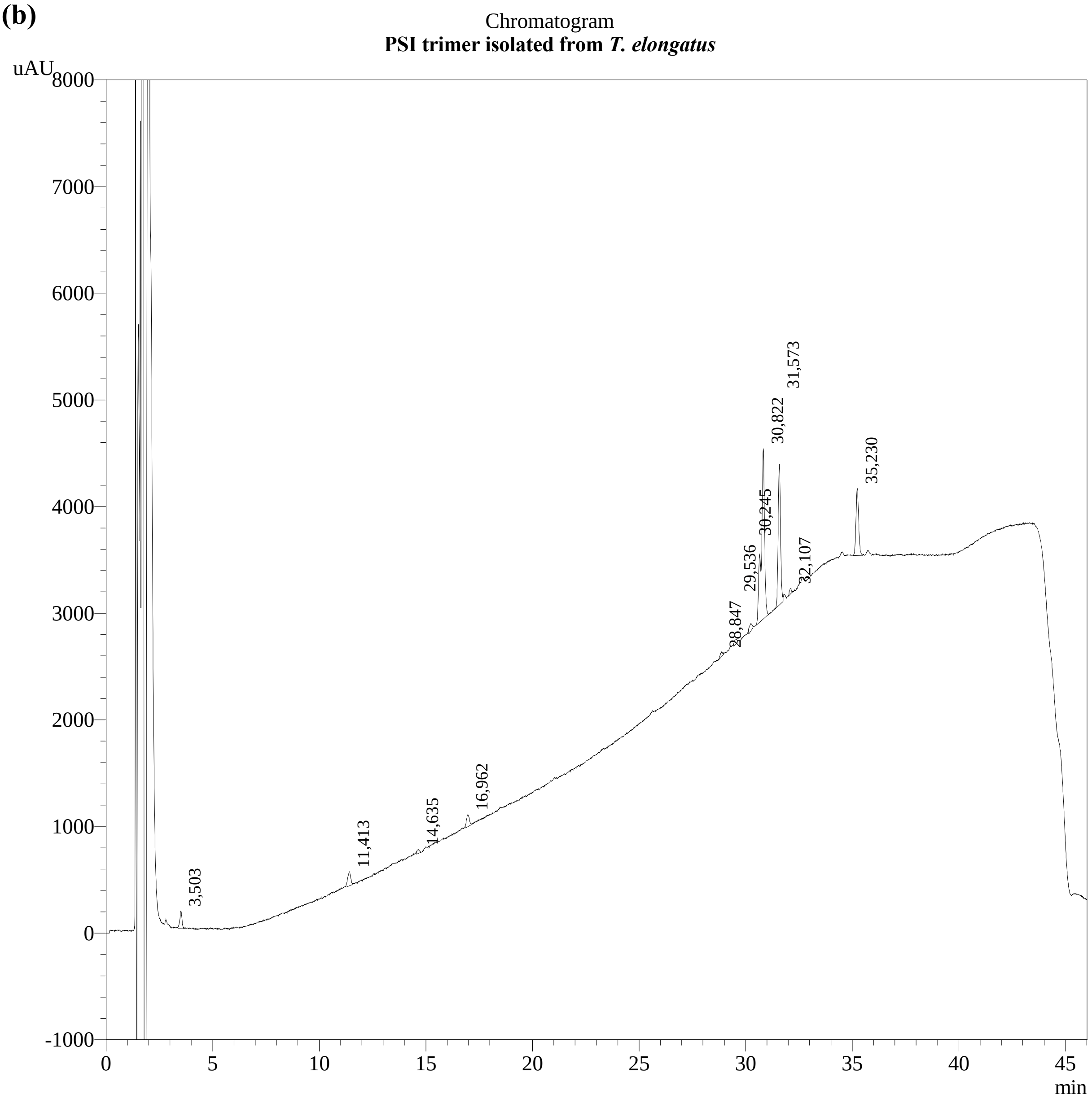


PeakTable

PDA Ch1 450nm

| Peak# | Name | Ret. Time | Area |
|---|---|---|---|
| 1 | unknown | 3,503 | 1195 |
| 2 | unknown | 11,413 | 1285 |
| 3 | unknown | 14,635 | 289 |
| 4 | unknown | 16,962 | 1046 |
| 8 | chla | 30,822 | 17733 |
| 9 | chla | 31,573 | 10340 |
| 10 | chla | 32,107 | 347 |
| 11 | beta carotene | 35,230 | 5388 |

**(c)**

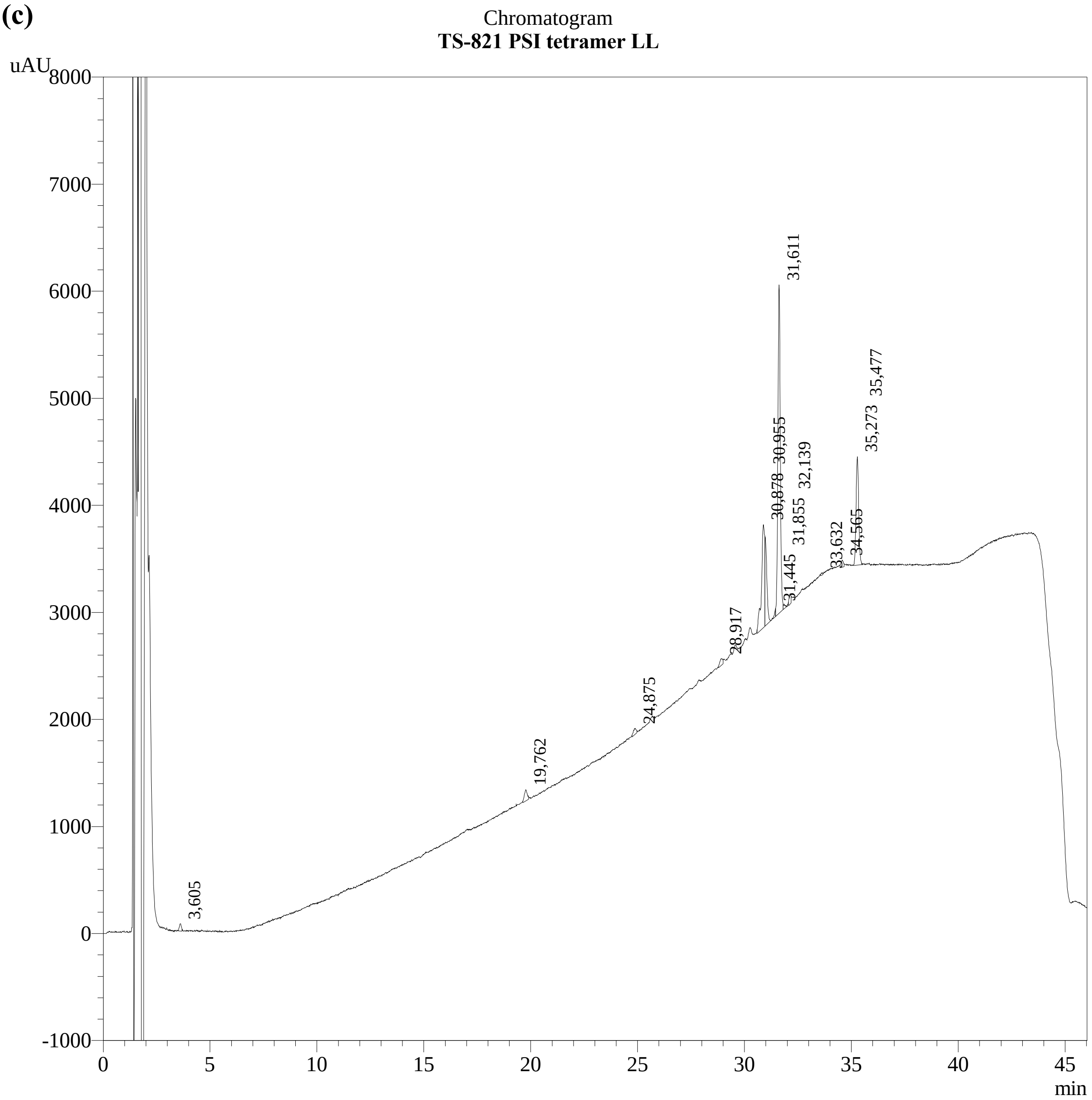


PeakTable

PDA Ch1 450nm

| Peak# | Name | Ret. Time | Area |
|---|---|---|---|
| 1 | unknown | 3,605 | 439 |
| 2 | myxo like | 19,762 | 985 |
| 3 | canta | 24,875 | 538 |
| 4 | unknown | 28,917 | 619 |
| 5 | chla | 30,878 | 10150 |
| 6 | echi | 30,955 | 4802 |
| 8 | chla | 31,611 | 24084 |
| 10 | chla | 32,139 | 578 |
| 11 | pht | 33,632 | 114 |
| 12 | unknown | 34,565 | 527 |
| 13 | beta carotene | 35,273 | 8588 |

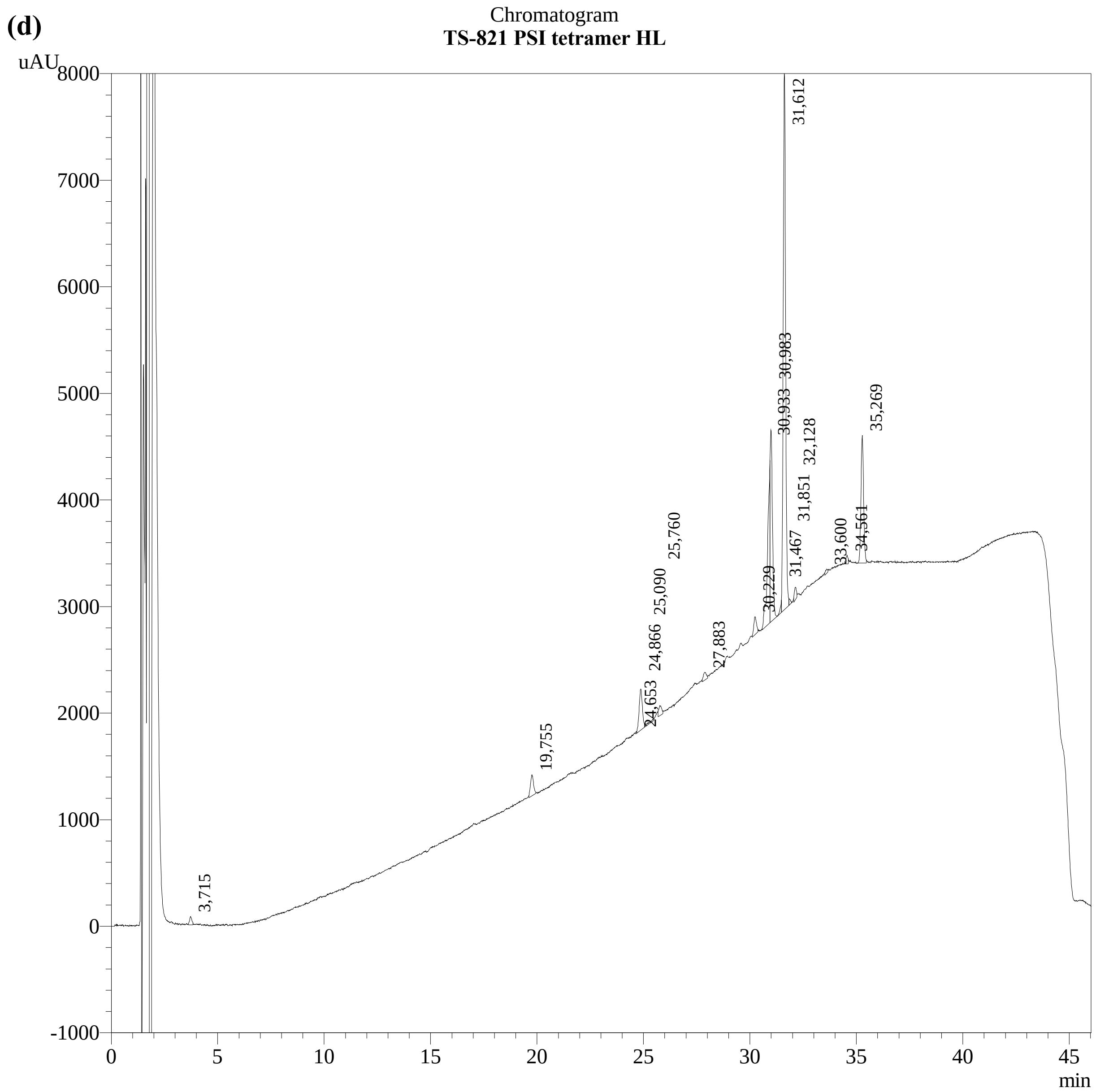


PeakTable

PDA Ch1 450nm

| Peak# | Name | Ret. Time | Area |
|---|---|---|---|
| 1 | unknown | 3,715 | 597 |
| 2 | myxo like | 19,755 | 1956 |
| 4 | canta | 24,866 | 4081 |
| 6 | unknown | 25,760 | 864 |
| 7 | unknown | 27,883 | 707 |
| 8 | unknown | 30,229 | 1493 |
| 9 | echi | 30,933 | 11826 |
| 10 | chla | 30,983 | 12822 |
| 12 | chla | 31,612 | 39687 |
| 14 | chla | 32,128 | 779 |
| 15 | pht | 33,600 | 289 |
| 16 | unknown | 34,561 | 488 |
|  | beta carotene | 35,269 | 10269 |

**Fig. S11. HPLC chromatograms of pigment analyses.**
The HPLC traces for pigment analyses of isolated inter-dimeric PSI carotenoids **(a)**, *T. elongatus* PSI trimer **(b)**, TS-821 PSI tetramer isolated from LL **(c)** and HL **(d)** are presented with their annotation tables under the chromatograms. Data acquisition and annotation were done by DHI (Denmark). *T. elongatus* PSI trimer has a Car/Chl ratio at 0.045 using this methodology.

**Table S1. PCR primers and conditions for different *psaL* cloning**

| Strain | Primers | Annealing |
|---|---|---|
| *Mastigocladus* sp. B 2965 | PsaF156/gmkrB1A | 43 °C |
| *Calothrix parietina* LB 1952 | PsaFr156/gmkB1C | 52.3 °C |
| *Scytonematopsis contorta* B 2963 | PsaF156/gmkB2 | 52 °C |
| *Spirirestis rafaelensis* B 2660 | PsaF156/gmkB2 | 52 °C |
| *Anabaena flos-aquae* LB 2558 | PsaF156/gmkB2 | 52 °C |
| *Nodularia spumigena* PCC 73104 | PsaF156/gmkB2 | 47 °C |
| *Cylindrospermum licheniforme* B 2014 | PsaF156/gmkB2 | 52 °C |
| *Anabaena inaequalis* B 381 | PsaF156/gmkB2 | 52 °C |
| *Tolypothrix distorta* B 424 | PsaFB3/gmkB2 | 54.6 °C |
| *Scytonema crispum* LB 1556 | PsaFB3/gmkrB2 | 57.5 °C |
| *Fischerella ambigua* 1903 | PsaFFisch/gmkFisch | 52 °C |
| *Nodularia harveyana* B 2093 | PsaFr156/gmkB2 | 52 °C |
| *Calothrix membranacea* B 379 | PsaFr156/gmkB2 | 52 °C |
| *Gloeotrichia ghosi* LB 1920 | PsaF156/gmkB2 | 52 °C |

**Table S2. Primer sequences used for *psaL* cloning**

| Primer name | Sequence 5' to 3' |
|---|---|
| PsaF156 | ATG MSW MGA TTG TT GCT TTG |
| PsaFB3 | TGG ATT GGY TGG GTM GG |
| PsaFFisch | TGC AAA CAA TCA AAA AGC ARG |
| PsaFr156 | TGG ATH GGY TGG GTH GG |
| gmkrB1A | GAR TSG GCR GAA TTT GC |
| gmkB1C | GAR TGG GCW GAR TTT GC |
| gmkB2 | GAR TGG GCV GAA TTT GCK GG |
| gmkrB2 | GAR TGG GCV GAA TTT GC |
| gmkFisch | AAA GGB ACT TTA ATG CGA TCG |
